# Induced ERBB response and standing FAK dependency nominate separable KRAS-combination hypotheses in pancreatic cancer

**DOI:** 10.64898/2026.07.09.737590

**Authors:** Jake Y. Chen, Ehsan Saghapour, Nikhil Kurmachalam, Geetanjali Oishe, Zhandos Sembay

## Abstract

Pancreatic ductal adenocarcinoma (PDAC) is driven by oncogenic KRAS in roughly 90% of cases, and KRAS-pathway inhibition has finally become clinically active. Durable benefit, however, will require identifying the adaptive and baseline vulnerabilities that shape response to KRAS inhibition. Two resistance mechanisms have been proposed separately in the literature — receptor-tyrosine-kinase bypass of KRAS, and dependence on the adhesion kinase FAK — but whether they are one target class or two, and which should partner a KRAS inhibitor, is unresolved. We integrate public perturbation, dependency, and survival data to nominate them as *mechanistically separable* candidate combination partners.

Two findings define the separation. First, KRAS loss increases ERBB2/3 receptor expression. This appeared in both an inducible genetic KRAS-extinction model and, independently, in five PDAC lines treated with pharmacological KRAS-G12C/D inhibitors, while MAPK output collapsed as expected. The signal was clearest for ERBB2 and in the genetic model; in the small pharmacological cohort the effect was modest and its confidence intervals crossed zero, so we treat ERBB2/3 up-regulation as a candidate adaptive response — ERBB2-dominant and ERBB3-compatible — not a proven resistance mechanism. Second, focal adhesion kinase (FAK/PTK2) is the top-ranked standing druggable dependency within the KRAS/Src/RTK/adhesion network we examined (essential in 58% of pancreatic lines), yet it is *not* induced by KRAS shutdown. FAK dependency is present at baseline and, in DepMap, is statistically independent of a line’s KRAS dependency (Spearman ρ = +0.05, n.s.) — a genuinely standing vulnerability rather than a KRAS-rebound effect. The candidate adaptive response and the standing dependency are not positively co-regulated across the perturbed lines (pooled Spearman ρ = −0.43, but n = 8 and n.s., so this cannot by itself establish independence); we therefore treat them as separable on mechanistic grounds — each nominated by different data and engaged by a different drug — rather than as statistically demonstrated independent programs.

A Src-centered signaling-landscape analysis associates patient prognosis with the coordinated invasion-and-RTK program these nodes organize, rather than with any single transcript; this program remains prognostic after adjustment for a conventional EMT/stromal signature, which does not (Src-neighborhood per–standard-deviation OS hazard ratio 1.9, p ≈ 3 × 10⁻⁵; EMT signature null on adjustment). Together these results motivate a concrete, testable hypothesis: that FAK inhibition (a standing dependency) and ERBB inhibition (a candidate induced adaptive response) are *separable* candidate partners for a KRAS inhibitor, best evaluated as distinct arms of a biomarker-stratified platform. They also clarify why single-agent Src inhibition — a non-oncogene dependency tested as monotherapy, without a KRAS backbone, in advanced rather than micro-metastatic disease — was not positioned to surface either mechanism. No protein-level, phospho-signaling, or combination-response validation is performed here; all findings are computational nominations that require experimental validation before any clinical inference.

**Highlights:**

- KRAS inhibition is associated with an induced ERBB2/3 up-regulation — ERBB2-dominant, ERBB3-compatible — directionally reproduced across genetic KRAS extinction and pharmacological KRAS-G12C/D inhibition; the pharmacological effect is modest and underpowered
- Genome-wide dependency nominates FAK — essential in 58% of pancreatic lines — as the top-ranked standing candidate co-target within the KRAS network; FAK dependency is present at baseline, statistically independent of KRAS dependency, and not co-regulated with the induced ERBB response
- Patient prognosis associates with a Src-organized invasion-and-RTK program, not with *SRC*, *KRAS*, or any single-gene transcript, and this program stays prognostic after adjustment for a conventional EMT/stromal signature
- The two mechanisms are separable on mechanistic grounds — nominated by different data and not positively co-regulated (though the direct correlation is underpowered, n = 8, n.s.) — motivating a multi-arm platform that could test FAK and ERBB partner arms as distinct hypotheses rather than one bundled combination

**In brief:** Blocking KRAS in pancreatic cancer is now clinically feasible, but resistance is the obstacle. Using only public data, Chen and colleagues nominate two mechanistically separable candidate combination partners for KRAS inhibitors: a candidate ERBB2-dominant adaptive (putative escape) response that is induced when KRAS is blocked, and FAK, the top-ranked standing dependency in the KRAS network — present at baseline and independent of a tumor’s KRAS dependency. Because the two are nominated by different data and are not positively co-regulated, they argue for a multi-arm KRAS-combination trial that tests each as a separate hypothesis — and they explain why the earlier single-agent Src trials, run without a KRAS backbone and in the wrong disease setting, were not positioned to detect either. The findings are computational nominations that require experimental validation.

## Introduction

Pancreatic ductal adenocarcinoma remains among the most lethal of common cancers, with five-year survival still in the low teens and a rising incidence that is projected to make it a leading cause of cancer death within the decade.^1^ The disease is genetically monotonous at its apex — oncogenic *KRAS* mutation is present in roughly 90% of tumors and, with *CDKN2A*, *TP53*, and *SMAD4* loss, defines the canonical progression.^2^ For most of that history KRAS was undruggable, and the field’s central problem was the absence of a way to attack the disease’s principal driver. That problem is now yielding: covalent G12C inhibitors are active in the small PDAC subset carrying that allele,^15^ and the multiselective RAS(ON) inhibitor daraxonrasib recently doubled overall survival in previously treated RAS-mutated pancreatic cancer.^14^ Alongside this, molecularly targeted options in PDAC remain confined to small biomarker-defined subsets, such as maintenance olaparib in germline *BRCA*-mutated disease,^16^ underscoring how few actionable dependencies the disease has offered. For the first time there is a backbone therapy against the driver — and, with it, a new and immediate question: what adaptive and baseline vulnerabilities does KRAS inhibition expose, and which can be targeted?

We approach that question with public data and nominate two candidate answers. The mechanistic literature has long placed receptor-tyrosine-kinase and stromal-bypass signaling at the center of RAS-pathway resistance.^10^ Oncogenic KRAS sustains a Src/PEAK1/ErbB2 amplification loop,^8^ Src-family kinases interact with KRAS and the YAP/TAZ axis to drive aggressive, drug-resistant disease,^9^ and cancer-associated-fibroblast–derived NRG1 enables ERBB-mediated bypass of KRAS inhibition.^10^ These circuits predict that removing KRAS should not simply kill the cell but reroute its signaling — and that the productive therapeutic targets are the nodes of that rerouting. Here we examine this prediction in perturbation data and in genome-wide dependency, and identify two candidate druggable nodes that behave differently: an *induced* ERBB2/3 up-regulation that appears when KRAS is shut off, and a *standing* FAK dependency that is present at baseline. We then relate patient prognosis to the coordinated invasion program these nodes organize, and argue that the earlier clinical failures of Src inhibition — the long-recognized target for exactly this invasion biology,^3^ including by AI-driven network analysis of metastatic PDAC^24^ — are consistent with a mis-deployment rather than a dead end. Throughout, we treat these as computational nominations: each is a hypothesis motivating experimental testing, not an established mechanism or a clinical recommendation.

This work is distinct from, and complementary to, the existing FAK-combination rationale in PDAC. Preclinical studies have shown that FAK inhibition remodels the immunosuppressive stroma and that combined FAK/RAF-MEK targeting reprograms fibroblasts,^12,13^ motivating trials such as RAMP-205 (avutometinib plus defactinib with chemotherapy). Our contribution is a synthesis and a separation: we rank FAK as the top standing dependency within the KRAS/Src/ RTK/adhesion network by genome-wide dependency data rather than treating it as one candidate among many; we show — with an independent replication cohort — that the receptors *induced* by KRAS shutdown are ERBB2/3, a response separate from FAK’s standing role and served by a different drug; and we therefore suggest that the two rationales are best tested as *separable* arms of a KRAS-anchored platform rather than bundled into a single combination whose result could not distinguish them. The novelty here is conceptual — the explicit separation of an induced adaptive-response axis from a standing-dependency axis — not a new molecular discovery, and it rests on public data rather than direct functional validation.

## Results

### KRAS inhibition is associated with an induced ERBB2/3 up-regulation

If KRAS-pathway inhibition is to be durable, the cell’s response to losing KRAS is where the next target is likely to be found. We asked what changes transcriptionally when KRAS signaling is removed, using two independent perturbation systems (Figure 2).

**Figure 1.**
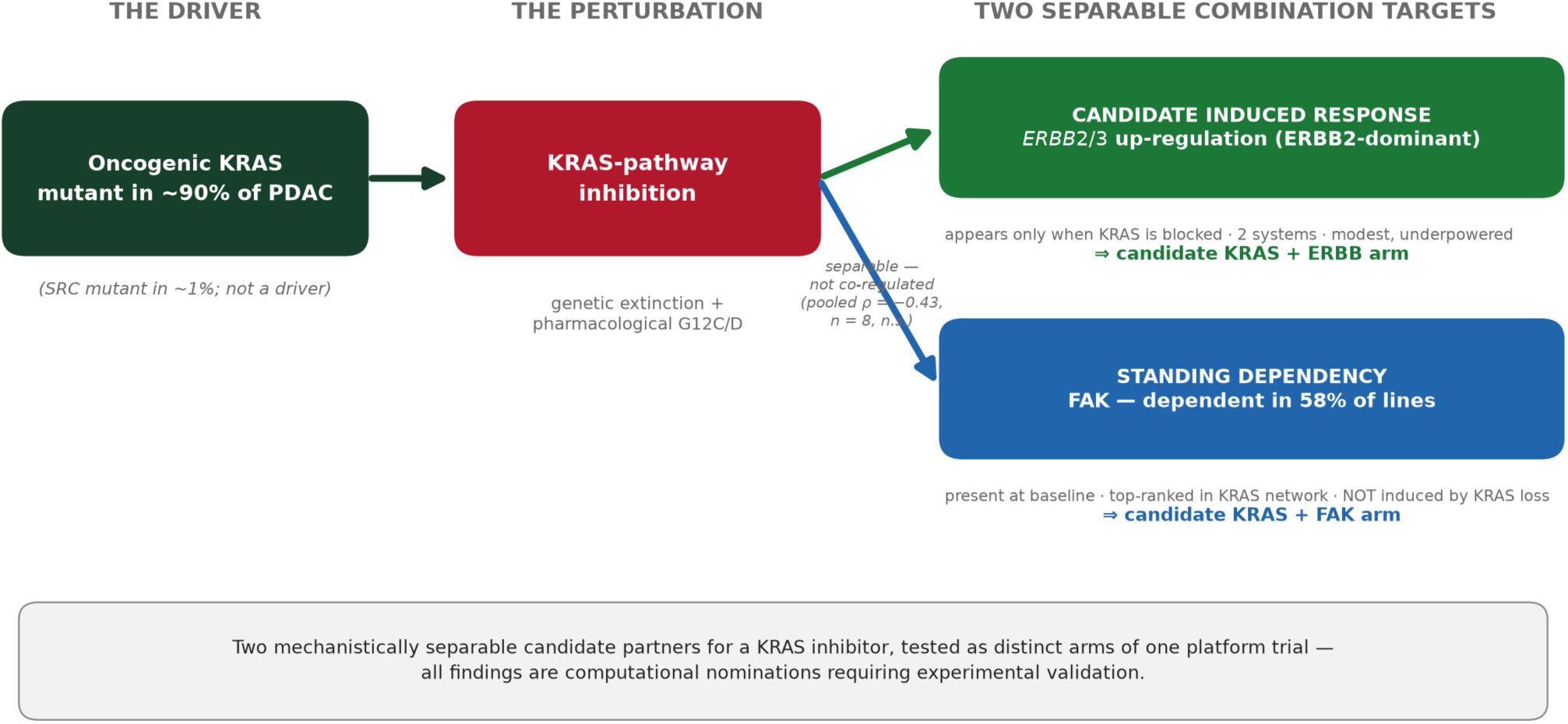
Two separable candidate combination targets for KRAS inhibition — study overview. A graphical summary: oncogenic KRAS (mutated in ∼90% of PDAC, Src in ∼1%) is inhibited; the cell responds by up-regulating an ERBB2/3 (and Src-family) receptor-tyrosine-kinase program — a candidate ERBB2-dominant adaptive (putative escape) response reproduced directionally across genetic and pharmacological KRAS inhibition in two independent datasets (modest, underpowered effect), while FAK — dependent in 58% of lines — is a standing dependency present at baseline and not induced. Because the two are nominated by different data and are not positively co-regulated, they motivate a multi-arm platform concept that would test KRAS + FAK and KRAS + ERBB as distinct partner arms. All findings are computational nominations requiring experimental validation.

**Figure 2.**
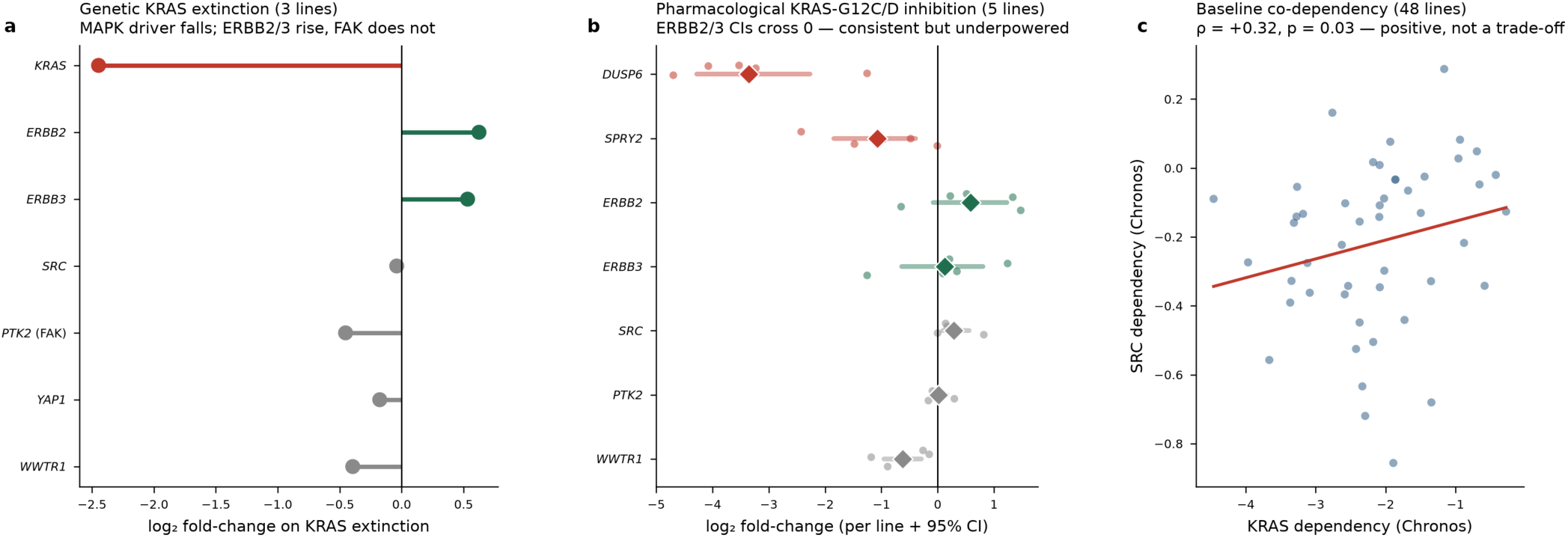
KRAS inhibition is associated with an induced ERBB2/3 up-regulation, directionally reproduced across two perturbation systems. (a) Genetic KRAS extinction across three inducible-KRAS PDAC lines (GSE240232): on KRAS shutdown, MAPK output collapses while ERBB2 (+0.62) and ERBB3 (+0.53) rise in 3/3 lines; FAK/PTK2 is not induced. (b) Pharmacological KRAS-G12C/D inhibition across five PDAC lines (GSE303051): the on-target genes DUSP6/SPRY2 indicate pathway shutdown, and the same candidate adaptive receptors ERBB2/3 rise in 4/5 lines (mean +0.58 and +0.13; bootstrap 95% CI [−0.08, 1.23] and [−0.64, 0.81] — both crossing zero, i.e. directionally consistent but not individually significant, p = 0.21 and 0.77), with Src-family SRC/FYN also rising while FAK is again not induced. Points are per-line log₂ fold-change across the five lines; bars the five-line mean. The per-line spread and the discordant line are shown to make the heterogeneity explicit. Across all eight perturbed lines, FAK response shows no positive coupling to ERBB2/3 induction (pooled Spearman ρ = −0.43, n.s.; ERBB2/3 up while FAK is not in 6/8 lines), so the induced up-regulation and the standing FAK dependency behave as separable programs. (c) Baseline co-dependency: across 48 pancreatic lines, SRC dependency correlates positively (not negatively) with KRAS dependency, indicating the inverse (escape) relationship is drug-induced rather than present at steady state. Asterisk, panel-level paired p < 0.05.

The first is an inducible genetic KRAS-extinction system across three PDAC lines: an engineered switch turns oncogenic KRAS off, and we measure which gene programs move. Shutting down KRAS collapses MAPK transcriptional output, as expected if the perturbation is on target (mean log₂ fold-change −1.80, paired p = 9×10⁻⁴), and *induces* the bypass receptors ERBB2 (+0.62, up in 3/3 lines, paired p = 0.007) and ERBB3 (+0.53, 3/3, p = 0.030); Src-family induction is modest (+0.15, 3/3, p = 0.021) and *SRC* itself is flat (−0.04, n.s.) (Figure 2a). This is the direction the KRas/Src/ErbB2 amplification loop^8^ and stromal-NRG1 ERBB bypass^10^ predict — but with only three lines at a single timepoint it is suggestive, not definitive.

We therefore looked for the same direction under an independent, *pharmacological* KRAS inhibition dataset: five PDAC cell lines (Pa01C, Pa03C, Pa14C, HPAC, MIA PaCa-2) treated with small-molecule KRAS-G12C/G12D inhibitors versus vehicle, in triplicate (GSE303051). The direction reproduces, though weakly (Figure 2b). On-target pathway shutdown is clear from the collapse of the direct MAPK-output genes *DUSP6* (−3.36, paired p = 0.005) and *SPRY2* (−1.07, p = 0.06). Against that background, ERBB2 rises in 4 of 5 lines (mean +0.58) and ERBB3 in 4 of 5 (+0.13), with the Src-family members *SRC* (+0.29, 4/5) and *FYN* (+0.41) also rising — the same directional pattern as the genetic system. We are explicit that these pharmacological effects are directionally consistent but statistically underpowered: the five-line paired tests do not reach significance for the candidate adaptive receptors (ERBB2 p ≈ 0.21; ERBB3 p ≈ 0.77), the per-line effect sizes are heterogeneous, and one line runs counter to the mean. The line-count summaries (“4/5 lines up”) describe consistency of direction, not strength of effect, and should be read that way. The two receptors are also not equally supported: ERBB2 shows the larger and more consistent induction in both systems (genetic +0.62, pharmacological +0.58), whereas ERBB3 is strong genetically (+0.53) but weak pharmacologically (+0.13, p ≈ 0.77). We therefore describe the induced adaptive response as **ERBB2-dominant and ERBB3-compatible**: ERBB2 is the better-supported node, and ERBB3 is consistent with the same direction but rests on thinner evidence. What can be said is that two mechanistically independent perturbations — genetic ablation and small-molecule inhibition — move the ERBB axis, and ERBB2 in particular, in the same direction; establishing that this transcript-level change is a functional escape will require protein/phospho-level and combination-perturbation experiments (see Discussion). By contrast, *PTK2* (FAK) is not induced in either system (pharmacological +0.01, 2/5; genetic −0.45, 0/3), which is the key negative that separates FAK from the induced response.

This induced pattern is invisible at baseline, which is why it took a perturbation to see it. Across the same 48 pancreatic lines, steady-state *SRC*, *YAP1*, and *WWTR1* dependency all correlate *positively* with KRAS dependency (Src Spearman ρ = +0.32, p = 0.03), not negatively as a standing trade-off would require (Figure 2c): at baseline Src and KRAS are co-dependent, and the inverse (escape) relationship appears only when KRAS is actively removed. The ERBB axis is further supported in patients independently of the perturbation: in two cohorts, *SRC* co-expresses with *ERBB2/3* — the receptors mediating CAF-NRG1 KRAS-bypass^10^ — and the correlation survives tumor-purity control (*ERBB3*: Moffitt ρ = +0.39, TCGA ρ = +0.59). The induced ERBB up-regulation is thus consistent across two orthogonal lines of evidence — dynamic perturbation in two systems, and static patient co-expression — while remaining, on the strength of the effect sizes alone, a nomination rather than a settled result.

### FAK is the top-ranked standing dependency in the KRAS network, and is also network-central

The induced ERBB response is one candidate target class; a second, mechanistically distinct candidate is present without any perturbation at all. Here we asked a different question from the escape analysis: not which genes rise after KRAS inhibition, but which genes PDAC cells already require for survival at baseline. We addressed it two ways — by functional CRISPR dependency and, orthogonally, by network centrality.

The functional criterion uses genome-wide CRISPR dependency from DepMap, where a more negative Chronos gene-effect score means that knocking out a gene more strongly impairs growth. Within the KRAS/Src/RTK/adhesion network we examined — a pre-specified set of driver, Src-family and RTK bypass nodes, and adhesion/invasion effectors — focal adhesion kinase (FAK/PTK2) is dependent (Chronos < −0.5) in 58% of pancreatic lines (bootstrap 95% CI 42–71%) and TAZ (WWTR1) in 38% (24–53%), against Src’s 13% (4–24%) and its paralogs YES1 (2%) and FYN (0%) (Figure 3a). FAK ranks second only to KRAS itself (93%, CI 84–100%). On the continuous Chronos scale, mean pancreatic-line gene effect is −0.61 for FAK versus −0.29 for Src, so FAK sits near the dependency boundary while Src remains weak throughout. All dependency percentages refer to the 45 pancreatic-adenocarcinoma lines with complete DepMap data; a 48-line subset used only for paired Src/KRAS bootstrap and co-dependency analysis is reported separately in STAR Methods. This rank is *within the examined KRAS-centered network*, not genome-wide: FAK is the top-ranked standing druggable node in this set relevant for KRAS-combination partnership, but is not claimed as the single strongest dependency genome-wide. FAK is prioritized over next-ranked nodes for reasons beyond percentage alone: TAZ/ WWTR1 (38%) is a transcriptional co-activator lacking selective clinical-stage inhibitors, and EGFR (36%) has failed in biomarker-unselected PDAC combinations; FAK has selective inhibitors in active PDAC trials and a mechanistic anti-stromal rationale. FAK is therefore best described as the *prioritized* standing co-target candidate — chosen on the joint basis of dependency rank, druggability, and translational package — not as a demonstrated post-KRAS-inhibition resistance dependency.

**Figure 3.**
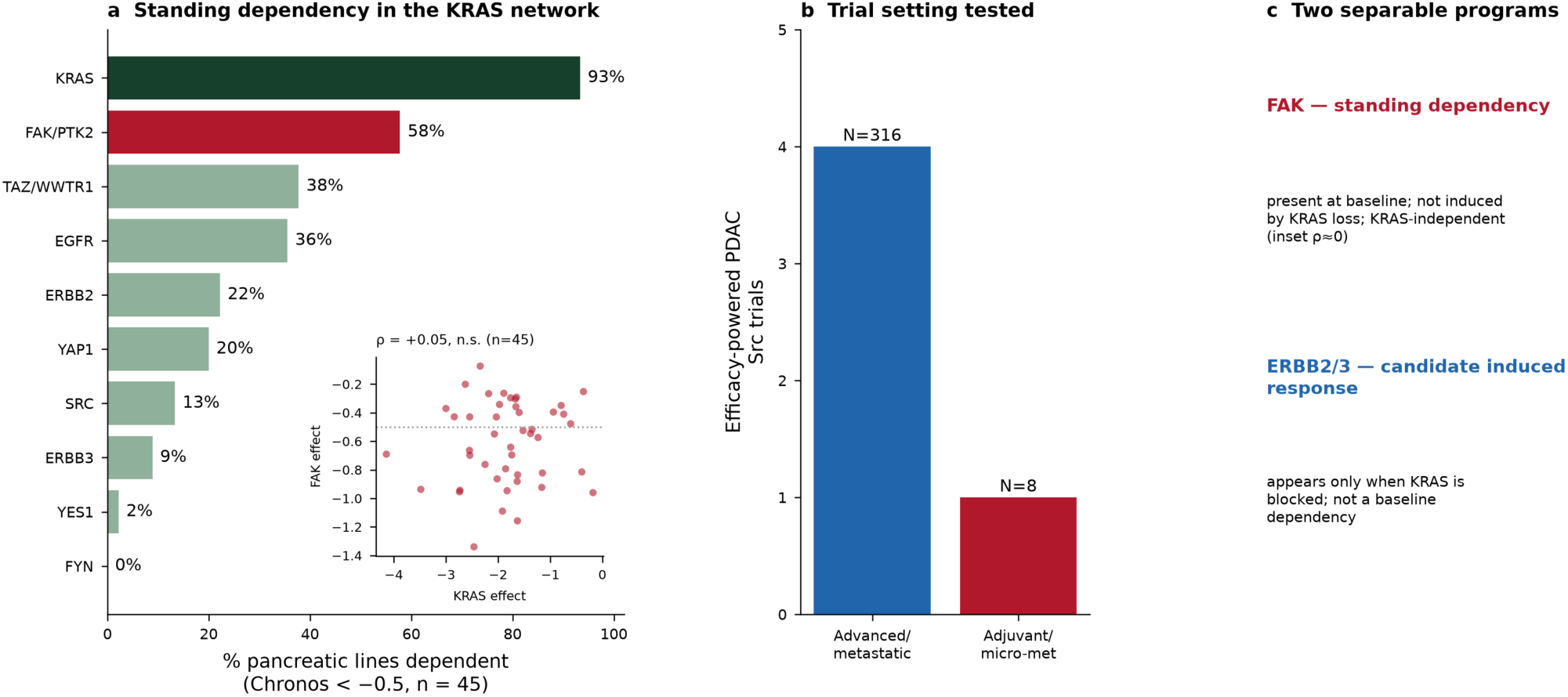
FAK is the top-ranked standing druggable dependency in the KRAS network, and the failed Src trials tested the wrong setting. (a) Fraction of the 45 pancreatic-adenocarcinoma cell lines with complete DepMap data that are dependent (Chronos < −0.5) across the KRAS network: KRAS 93%, FAK/PTK2 58%, TAZ/WWTR1 38%, EGFR 36%, ERBB2 22%, YAP1 20%, SRC 13%, ERBB3 9%, YES1 2%, FYN 0%. FAK is the top-ranked node after KRAS and is distinct from the candidate induced ERBB response; a WINNER network-centrality ranking within this pre-specified module (not a genome-wide search) places FAK 4th of 50 (ranking p = 0.0016 vs a 10,000-network degree-preserving null), corroborating the functional dependency within the curated candidate set. Inset (panel a): FAK/PTK2 gene effect is uncorrelated with KRAS gene effect across 45 PDAC lines (Spearman ρ = +0.05, n.s.; dotted line, dependency threshold), indicating a standing rather than KRAS-linked vulnerability. (b) Efficacy-powered PDAC Src trials by disease setting: four (N = 316) tested macroscopic advanced/metastatic disease with cytoreduction endpoints, only one (N = 8) the resected/adjuvant micro-metastatic setting where Src’s invasion biology is predicted to matter. (c) The two nominated programs are separable: FAK is a baseline standing dependency (KRAS-independent), whereas ERBB2/3 up-regulation appears only under KRAS inhibition.

The network criterion is independent of dependency data and uses the authors’ WINNER algorithm^25^ — a personalized-PageRank node-prioritization over a weighted protein-interaction graph, with significance assessed against 10,000 degree-preserving random networks. Seeding the same KRAS/Src/RTK/adhesion module on a STRING-derived interactome (50-gene ranked universe), FAK/PTK2 ranks 4th of 50 by network centrality, unlikely under the degree-matched null (ranking p = 0.0016); Src ranks 1st and EGFR 3rd, while broad-composite and low-connectivity nodes fall lower (Figure 3a, inset; Supplementary Figure S-SPINNER and Supplementary WINNER table). The full prioritized interactome — node size scaled to WINNER centrality, nodes coloured by rank significance against the null, and edges drawn from the STRING confidence graph in the SPINNER application — is shown in Supplementary Figure S-SPINNER; notably FN1 ranks 2nd by raw centrality but is not significant against the degree-matched null (p = 0.95), whereas FAK’s rank-4 position is (p = 0.0016), which is why connectivity alone does not nominate a target. The two criteria are complementary: network centrality reflects interactome connectivity (on which Src ranks highest), whereas CRISPR dependency reflects functional survival requirement (on which FAK ranks highest after KRAS). FAK is distinctive in scoring near the top on *both* — a highly-central node (rank 4, p = 0.0016) that is also the strongest standing functional dependency — whereas Src is central but not dependent and broad-composite nodes are neither. That convergence, by different mathematics on different data, elevates FAK from one candidate to the prioritized standing co-target.

WINNER re-ranks *within* the pre-specified KRAS/Src/RTK/adhesion module and its STRING neighborhood, so it is not unbiased genome-wide discovery and is not independent biological validation: it shows that among genes already chosen for KRAS-network relevance, FAK is unusually central given its connectivity. The centrality analysis is therefore a prioritization consistency check on a curated candidate set, not independently-arising evidence. The functional CRISPR dependency is the stronger criterion — it draws on genome-wide DepMap data — while network centrality corroborates it within that module. A fully unbiased test would rank FAK against the entire kinome or druggable genome, which we have not done.

Because a KRAS-combination partner should ideally be a vulnerability the tumor carries regardless of KRAS state, we tested whether FAK dependency is KRAS-linked. Across 45 PDAC lines, FAK (*PTK2*) gene effect does not correlate with KRAS gene effect (Spearman ρ = +0.05, p = 0.75; bootstrap 95% CI −0.25 to +0.35), and the fraction of FAK-dependent lines is essentially identical in the more-KRAS-dependent and less-KRAS-dependent halves (57% versus 59%; Mann–Whitney p = 0.85) (Figure 3a, inset). FAK dependency is therefore a *standing* vulnerability, present across the cohort independent of KRAS state, rather than a KRAS-rebound effect. This contrasts with the ERBB result: ERBB2/3 matters only once KRAS is removed, whereas FAK matters at baseline. One boundary on this claim should be stated plainly: what we test is independence from baseline KRAS *dependency* (CRISPR gene-effect magnitude), not from KRAS *inhibition*. FAK is not transcriptionally induced by KRAS shutdown (Figure 2a), but whether its functional essentiality changes under pharmacological KRAS blockade — a drug-anchored FAK-dependency measurement — has not been tested here and is part of the experimental package a trial would require.

FAK also carries the deepest translational package. FAK inhibition reduces desmoplasia and restores CD8 T-cell infiltration in genetically engineered pancreatic models,^12^ and combined FAK plus RAF/MEK targeting reprograms cancer-associated fibroblasts toward regression and improves responses to cytotoxic and immune therapy^13^ — the basis of the active RAMP-205 trial. The ranking nominates FAK as the top-ranked *standing* dependency in the KRAS network and, given the anti-stromal preclinical package, as a rational KRAS-combination partner on a constitutive-vulnerability rationale. It does not show that FAK is specifically required to survive the *post-KRAS-inhibition* state — FAK is not induced by KRAS shutdown, so its rationale is standing vulnerability, not escape-handling. This is why FAK and ERBB are best tested as distinct arms: they are nominated by different evidence (baseline dependency versus perturbation induction), are not co-regulated, and are drugged by different agents. Across eight perturbed lines with per-line data, the magnitude of FAK (*PTK2*) response on KRAS shutdown is not positively correlated with ERBB2/3 induction — the point estimate is weakly *negative* (pooled Pearson r = −0.33; Spearman ρ = −0.43), though with only eight lines this is not significant (p = 0.29). In six of eight lines ERBB2/3 rises while FAK does not (FAK mean log₂FC −0.16, up in only 2/8) — FAK a baseline vulnerability not amplified by KRAS shutdown, ERBB2/3 specifically induced. That separability is the conceptual contribution and the design principle of the platform outlined in the Discussion.

### The failed Src trials tested the wrong setting, not a worthless target

The kinase historically nominated for this adhesion-and-invasion biology is Src, and its clinical record is uniformly negative — a fact that has been read as closing the target. That reading conflates the target with its deployment. Src’s reported biology in PDAC is invasion, epithelial-mesenchymal plasticity, adhesion turnover, and stromal crosstalk^4,5,11^ — biology that would be expected to matter against minimal residual disease rather than in the shrinkage of established macroscopic tumors. The trials did the opposite (Figure 3b). Of the five efficacy-powered PDAC Src trials, four (total N = 316) tested macroscopic advanced or metastatic disease with cytoreduction endpoints, the setting where a non-survival dependency is least likely to help; only one (N = 8) tested the resected, adjuvant, micro-metastatic setting, far too small to interpret. The pivotal negatives are real — dasatinib plus gemcitabine gave no survival benefit in locally advanced disease (HR 1.16, 95% CI 0.81–1.65, p = 0.57)^6^ and saracatinib plus gemcitabine did not improve over gemcitabine alone^7^ — but both, and all the others, were single-agent additions run without a KRAS inhibitor, so neither the candidate induced adaptive response nor the invasion setting in which Src biology is proposed to operate was positioned to be surfaced. We present this as a reinterpretation consistent with the data, not as proof that a correctly-deployed Src strategy would succeed.

### Patient prognosis associates with a Src-organized invasion-and-RTK program, not with SRC, KRAS, or any single transcript

To relate these mechanisms to patient outcome, we turn to a prognostic analysis that we flag at the outset as the most exploratory in the paper: it is single-cohort, uses an investigator-curated gene module and an in-house pipeline, and is presented as hypothesis-generating context requiring external validation, not as an independent prognostic claim. With that caveat foremost, we used GeneTerrain — a method that lays a tumor’s expression profile onto a fixed two-dimensional “map” of a gene module, so that programs of co-regulated genes appear as regions of the map rather than as a list; a summary score then measures how active each region is. We applied it to TCGA-PAAD (178 primary tumors, with four solid-normal samples as the baseline; Figure 4). A 42-gene SRC-centered module spanning five programs (SRC-pathway core, EMT/ invasion, CAF/fibrosis, immune evasion, epithelial identity) tracks the aggressive PDAC state: tumors high on composite SRC-terrain activity are enriched for invasion and stromal machinery — ZEB1/2, VIM, FN1, SNAI2 (EMT); COL1A1/2, COL3A1, FAP, PDGFRB (CAF/fibrosis); MMP2, MMP14 (matrix remodeling) (Supplementary Figure S-GeneTerrain).

**Figure 4.**
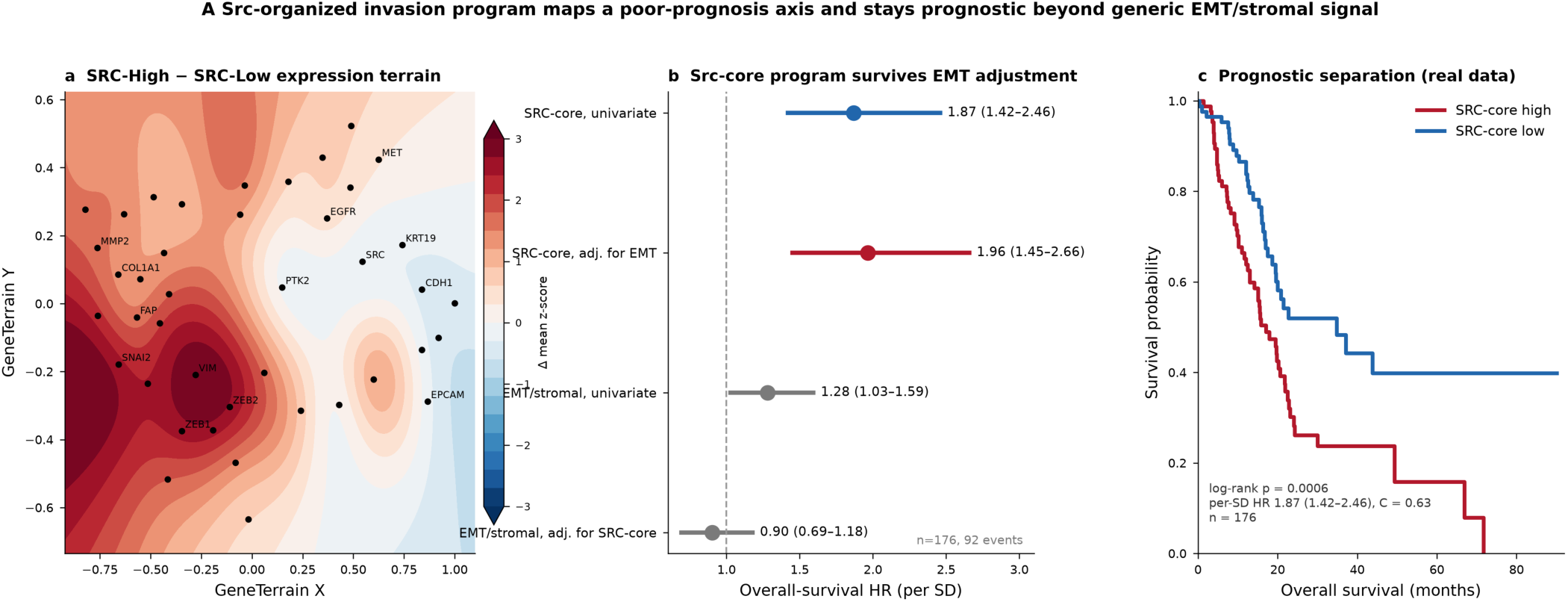
A Src-organized invasion program maps a poor-prognosis axis and stays prognostic beyond a generic EMT/stromal signature (TCGA-PAAD; exploratory). (a) GeneTerrain difference map: each of the 42 module genes is placed at a fixed 2-D coordinate, and colour shows the mean normal-baseline z-score difference between SRC-High and SRC-Low tumours (top vs bottom quartile of composite SRC activity). The invasion/EMT/CAF cluster — SNAI2, ZEB1/2, VIM, FN1, the collagens, FAP, MMP2 — is coordinately elevated in SRC-High tumours (warm), while epithelial-identity genes (EPCAM, CDH1, the keratins) are depleted (cool), so the aggressive state appears as a coherent region of the map rather than a scattered gene list. (b) Per-SD overall-survival hazard ratios for a Src-signaling-neighborhood score and a conventional EMT/ stromal signature, univariate and mutually adjusted (n = 176, 92 events). Both are prognostic alone, but in the joint model the Src-neighborhood score retains its full effect (HR 1.96, 95% CI 1.45–2.66) while the EMT signature collapses to null (HR 0.90) — the prognostic information is not reducible to generic mesenchymal/stromal content. (c) Kaplan–Meier overall survival for patients split at the median Src-neighborhood score (real per-patient TCGA-PAAD data; median-split log-rank p = 6×10⁻⁴; per-SD OS HR 1.87, 95% CI 1.42–2.46; concordance 0.63; n = 176). The full multi-panel GeneTerrain dashboard — network topology, per-group terrain surfaces, module-level separation, and per-gene survival associations — is provided as Supplementary Figure S-GeneTerrain, and the entire analysis is reproducible from public data with a standalone harness (Supplementary Data; no dependence on GeneTerrain as a service). The EMT-adjustment confounder check uses a 10-gene Src-neighborhood proxy, not the 42-gene module itself; the association is single-cohort and exploratory and requires external validation.

Within this cohort, outcome associates with the program rather than the single gene. The cleanest prognostic signal is the SRC-pathway *core* program: per-score overall-survival HR 1.96 (95% CI 1.44–2.65, p = 1.5×10⁻5), equivalent to a per–standard-deviation OS HR of 2.24 and PFI HR of 2.01, with concordant Kaplan-Meier separation (log-rank p = 0.011 OS, 0.0036 PFI). A coherent poor-prognosis axis of receptor-tyrosine-kinase, adhesion, matrix-invasion, and EMT effectors — MET, KRT19, ITGA2, VCL, EGFR, MMP14, MMP7, SNAI2, each surviving BH-FDR correction — underlies it. The broad composite terrain score, by contrast, is only weakly and non-robustly associated with survival (per-score HR 1.31, 95% CI 1.02–1.68, p = 0.037). The full panels are in Supplementary Figure S-GeneTerrain; the main Figure 4 distills the one result that matters for this paper — that the Src-organized program stays prognostic after adjustment for a conventional EMT/stromal signature (per-SD HR 1.96 joint, versus EMT null; Figure 4b), so the outcome association concentrates in the coordinated invasion-and-RTK program, the same adhesion/ERBB biology the perturbation and dependency analyses point to, not in any single transcript. As noted above, the program-level association may in part reflect stromal content, tumor purity, or molecular subtype and requires external validation across independent cohorts before it can be considered more than corroborative; the EMT-adjustment check uses a 10-gene proxy, not the 42-gene module itself. Because a prior concern with this analysis was its reliance on an in-house pipeline, we note that the entire GeneTerrain result is now reproducible from public TCGA-PAAD data by a standalone, dependency-listed harness — with checksummed inputs and outputs — that does not call GeneTerrain as a service (Supplementary Data). Re-running it reproduces the SRC-pathway–core prognostic signal (OS HR 2.24, p = 1.3×10⁻5; PFI HR 2.01, p = 2.5×10⁻5) and the FDR-significant per-gene axis (Supplementary Figure S-GeneTerrain-repro).

As a first check on the most immediate confounder — whether the association is simply generic EMT/stromal biology — we asked whether a Src-signaling-neighborhood score adds prognostic information beyond a conventional mesenchymal/CAF signature in the same cohort. This is a proxy analysis, not a re-run of the 42-gene GeneTerrain pipeline: from the TCGA-PAAD expression matrix we built a 10-gene Src-neighborhood score (SRC, PTK2, EGFR, MET, ITGB1, ITGA2, VCL, MMP2, MMP14, KRT19) and a 14-gene EMT/stromal signature (mesenchymal and CAF markers), and compared them in Cox models over 176 tumors (92 events). Alone, both are prognostic (Src-neighborhood per-SD OS HR 1.87, 95% CI 1.42–2.46, p = 8×10⁻⁶; EMT HR 1.28, 1.03–1.59, p = 0.03), but the two are only moderately correlated (r = 0.58), and in a joint model the Src-neighborhood score retains its full effect (HR 1.96, 1.45–2.66, p = 1×10⁻5) while the EMT signature collapses to null (HR 0.90, p = 0.45); concordance is 0.63 for the Src-neighborhood score versus 0.51 for EMT. The prognostic signal in this Src-organized program is therefore not reducible to generic EMT/stromal content — it carries information a simple mesenchymal signature does not. This supports, but does not by itself validate, the GeneTerrain observation; a full external replication of the 42-gene module in an independent cohort remains required.

### Src has the profile of a combination-modality target, not a monotherapy

The two mechanisms above place Src within a combination rather than as a single agent, and its molecular profile explains why (Figure 5). Src is neither a genetic driver (*SRC* mutated in 0–1.4% of PDAC versus *KRAS* 62–96%; Figure 5a) nor a survival dependency (genetic *KRAS* ablation lethal in 83% of the 48 lines, *SRC* in none; Figure 5b,c), and it is broadly expressed in normal tissue (Figure 5d) — leaving no *SRC* genotype for selection and little window for a systemic single agent. Its reported “high Src, worse survival” association (HR ≈ 1.45)^24^ does not hold as a continuous effect (pooled per-SD HR 0.89, 95% CI 0.69–1.14; Figure 6) and appears only under a p-minimizing cutpoint (permutation-corrected p = 0.12; full analyses in Supplementary Methods §6). Src is therefore best regarded as one node within the KRAS-combination biology above — ideally engaged through a scaffold-blocking or degrading agent rather than a catalytic inhibitor given alone — a secondary, supporting analysis included because the claim is in the literature and the authors’ prior work,^24^ not the paper’s central finding.

**Figure 5.**
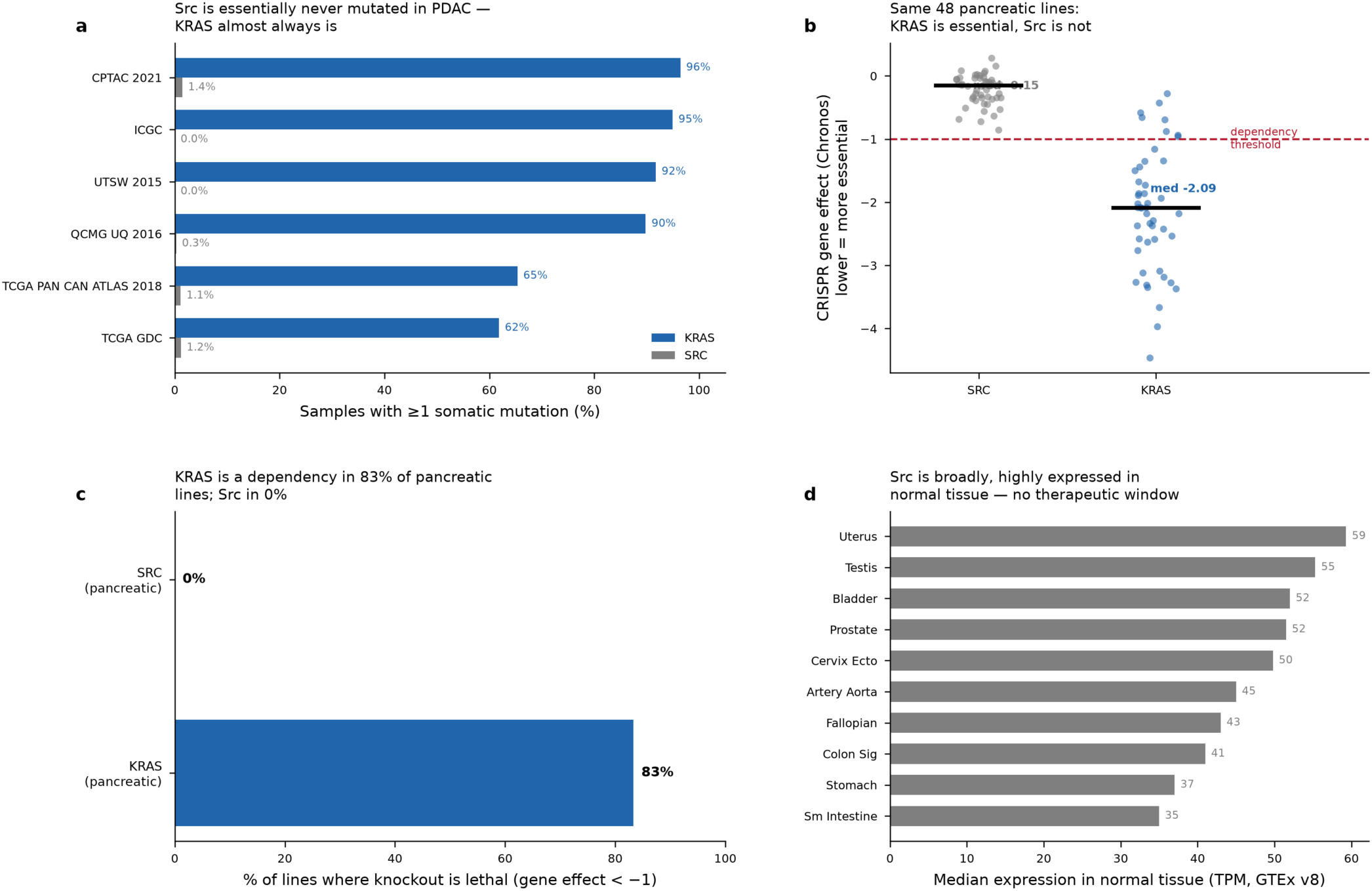
Src is a non-oncogene dependency, which is why single-agent inhibition cannot shrink a tumor. (a) Somatic mutation frequency of SRC (0–1.4%) versus KRAS (62–96%) across six PDAC cohorts. (b) Paired CRISPR gene effect for SRC and KRAS across 48 pancreatic lines; KRAS clusters below the dependency threshold, Src near zero. (c) Fraction of lines where knockout is lethal (Chronos < −1): KRAS 83%, SRC 0%. (d) SRC median expression across normal tissues (GTEx) — broad and high, leaving no therapeutic window for a systemic naked inhibitor.

**Figure 6.**
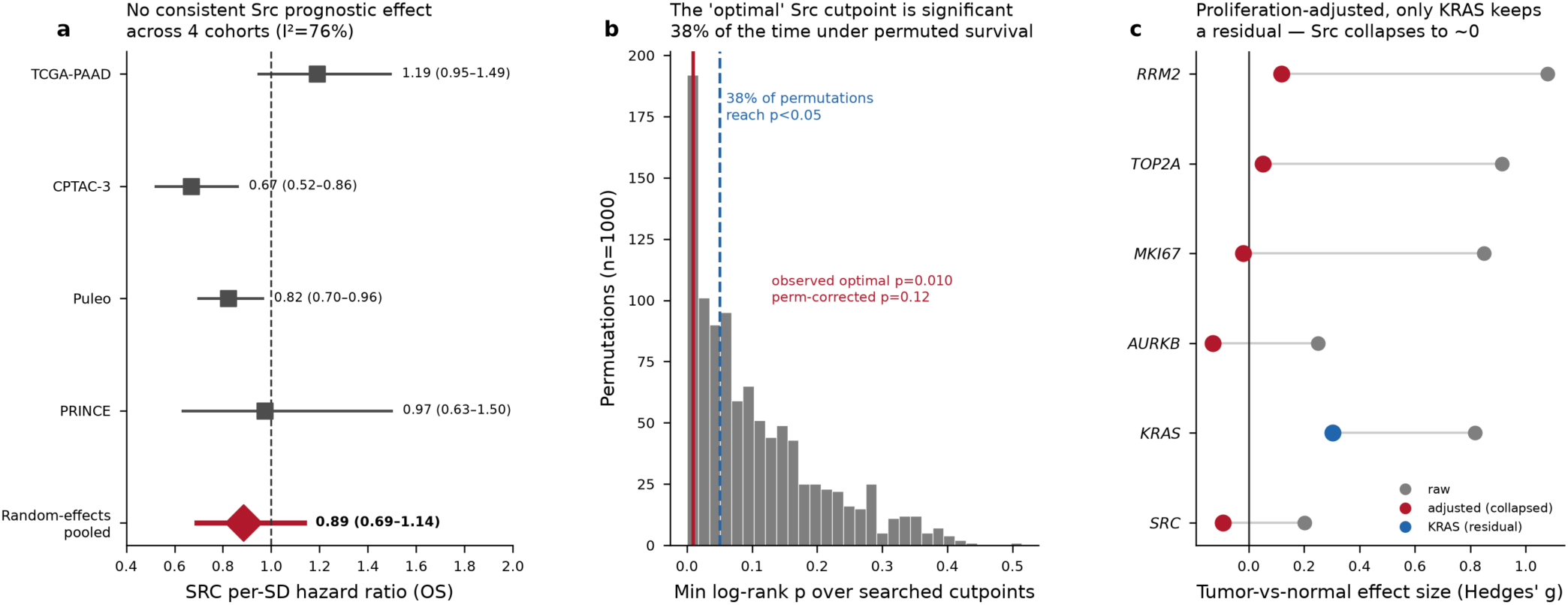
The reported Src prognostic signal is a cutpoint and proliferation artifact. (a) Continuous per-SD Src overall-survival hazard ratios across four cohorts (TCGA-PAAD, CPTAC-3, Puleo, PRINCE) with the random-effects pooled estimate: Src is never significantly hazardous (pooled HR 0.89, 95% CI 0.69–1.14; I² = 75.6%). (b) Permutation null of the minimum log-rank p over searched cutpoints in TCGA-PAAD; the “optimal” cutpoint reaches p < 0.05 in 38% of permutations, giving the observed optimum (HR 2.86, nominal p = 0.010) a permutation-corrected p of 0.12. (c) Leave-one-out proliferation adjustment in the Moffitt cohort: RRM2, TOP2A, MKI67, and Src collapse toward zero, while only KRAS retains a significant residual.

Two observations here may appear to conflict but do not, because they measure different objects. The single *SRC* transcript is not a robust prognostic marker (continuous per-SD HR ≈ 0.9; the HR ≈ 1.45 recovers only under an optimized cutpoint), yet the Src-organized invasion program is strongly associated with survival (per-SD HR ≈ 1.9, p ≈ 10⁻5) — the expected result if the prognostic biology is distributed across a module rather than carried by one probe. The reinterpretation therefore refines rather than negates the prior single-gene nomination: the actionable signal lives at the level of the invasion program and its most-dependent node (FAK), not the single *SRC* transcript.

### Two separable candidate combination classes

Taken together, the analyses nominate two mechanistically separable candidate partner classes for a KRAS inhibitor in PDAC. The first is an *induced* ERBB2/3 up-regulation that appears when KRAS is blocked — directionally reproduced across genetic and pharmacological perturbation, corroborated by patient co-expression, but of modest and statistically underpowered effect size. The second is a *standing* FAK dependency present at baseline — after KRAS itself, the top functional CRISPR dependency among the KRAS/Src/RTK/adhesion genes we examined, drawn from genome-wide DepMap gene-effect data and corroborated by high network centrality within the same curated module, and the most translationally advanced anti-stromal node in the disease. Because the two are nominated by different data, are not positively co-regulated, and are drugged by different agents, they are best evaluated as separate hypotheses rather than bundled into one combination whose result could not distinguish them. How that separation could be tested clinically — and what would first be required to justify a trial at all — is the subject of the Discussion.

## Discussion

KRAS inhibition has become clinically active in pancreatic cancer, and the next gain is likely to come from targeting the adaptive and baseline vulnerabilities it exposes. Using only public data, this study nominates two candidate routes and the targets that might address them. The first is an *induced* ERBB2/3 up-regulation, directionally reproduced across two mechanistically independent perturbations — genetic KRAS extinction and pharmacological KRAS-G12C/D inhibition — and corroborated by patient co-expression, though with effect sizes that are modest and, in the pharmacological cohort, not individually significant. The second is a *standing* FAK dependency, the top-ranked druggable node in the KRAS network by functional dependency and corroborated by network centrality within the same module, and the most translationally advanced anti-stromal target in the disease. That the two are separable — nominated by different data, not positively co-regulated, drugged by different agents (a mechanistic rather than a statistically-demonstrated separation) — is the conceptual contribution of the paper: it argues that a rational combination program should keep them as distinct hypotheses rather than merging them.

This offers a productive reframing of the long-standing Src question. The uniformly negative Src-inhibitor trials are better read not as a verdict on the underlying adhesion-and-invasion biology but as the result of a specific deployment — a non-oncogene dependency given as single-agent catalytic inhibition, for tumor shrinkage, in advanced macroscopic disease, without a KRAS backbone. Each of those choices is now correctable in principle. Within the cohort we examined, patient outcome associates with the Src-organized invasion-and-RTK program even though it does not associate with *SRC* transcript abundance. The productive question is therefore not whether a Src inhibitor shrinks a pancreatic tumor — it does not — but whether blocking the adhesion/RTK adaptive-response nodes delays relapse and resistance when combined with KRAS inhibition, in the setting where dissemination rather than bulk is the concern. This remains a hypothesis, not a demonstrated effect.

### A candidate trial design, and what would have to precede it

The following is speculative scaffolding, contingent on the Box 1 validation package returning positive; it is offered to make the separability principle concrete, not as a design we advocate on current evidence. Should the nominations survive that validation, the separability principle would suggest a specific clinical test: a biomarker-stratified, multi-arm platform anchored on a KRAS inhibitor — KRAS inhibitor alone (shared control), KRAS + FAK inhibitor (the standing-dependency, anti-stromal rationale), and KRAS + ERBB inhibitor (the induced adaptive-response rationale) — so that each mechanism is tested against one control rather than bundled (Figure 7). A workable frame would take progression-free survival as the primary endpoint in the advanced/KRAS-inhibitor–experienced setting, with each partner arm compared against the shared control and a moderate hazard-ratio target; the adjuvant/minimal-residual-disease variant would use disease-free survival with a cure-fraction analysis. (Formal power, α-allocation, and per-arm sample sizes are deferred to a full protocol, since they depend on the validation read-outs above.) The projected curves in Figure 7 are illustrative of this structure only — Weibull models with the control arm anchored to the RASolute 302 daraxonrasib benchmark (OS 13.2 vs 6.7 months, HR 0.40)^14^ and partner arms drawn under an assumed additive effect — and depict trial geometry, not predicted efficacy. We emphasize that these numbers are design scaffolding, not a protocol, and that no clinical recommendation follows from this work. Candidate agents by class are summarized in Table 1.

**Figure 7.**
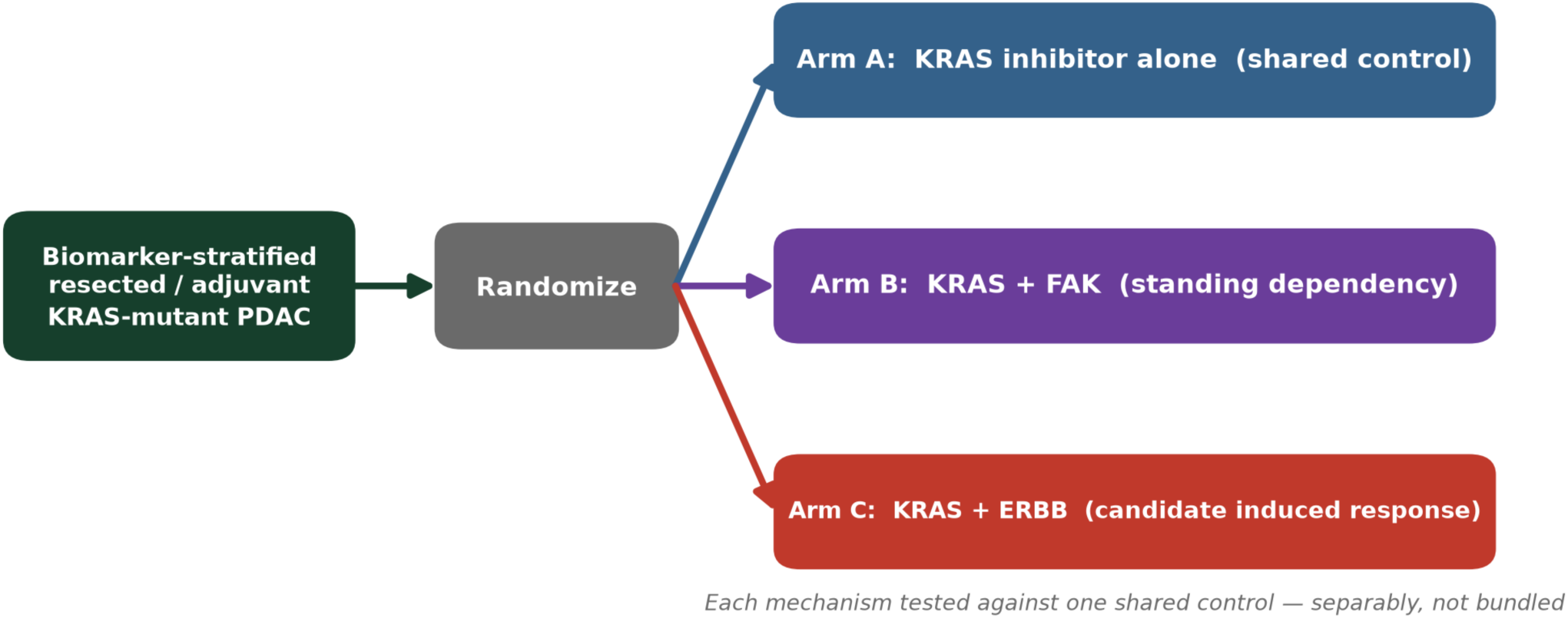
An experimental-to-clinical translation framework, not a protocol. A schematic of how the two nominations *would* be tested separably if they survive the validation experiments of Box 1: biomarker-stratified resected/ adjuvant KRAS-mutant PDAC randomized to a KRAS inhibitor alone (Arm A, shared control), KRAS + FAK (Arm B, standing dependency), or KRAS + ERBB (Arm C, candidate induced response), so each rationale is tested against one common control rather than bundled into a single combination whose result could not attribute benefit. This is a conceptual design framework illustrating attribution logic, not a trial protocol or a recommendation for current practice; no efficacy is implied. (Illustrative projected-geometry curves are provided only in Supplementary Figure S-proj.)

**Table 1.**
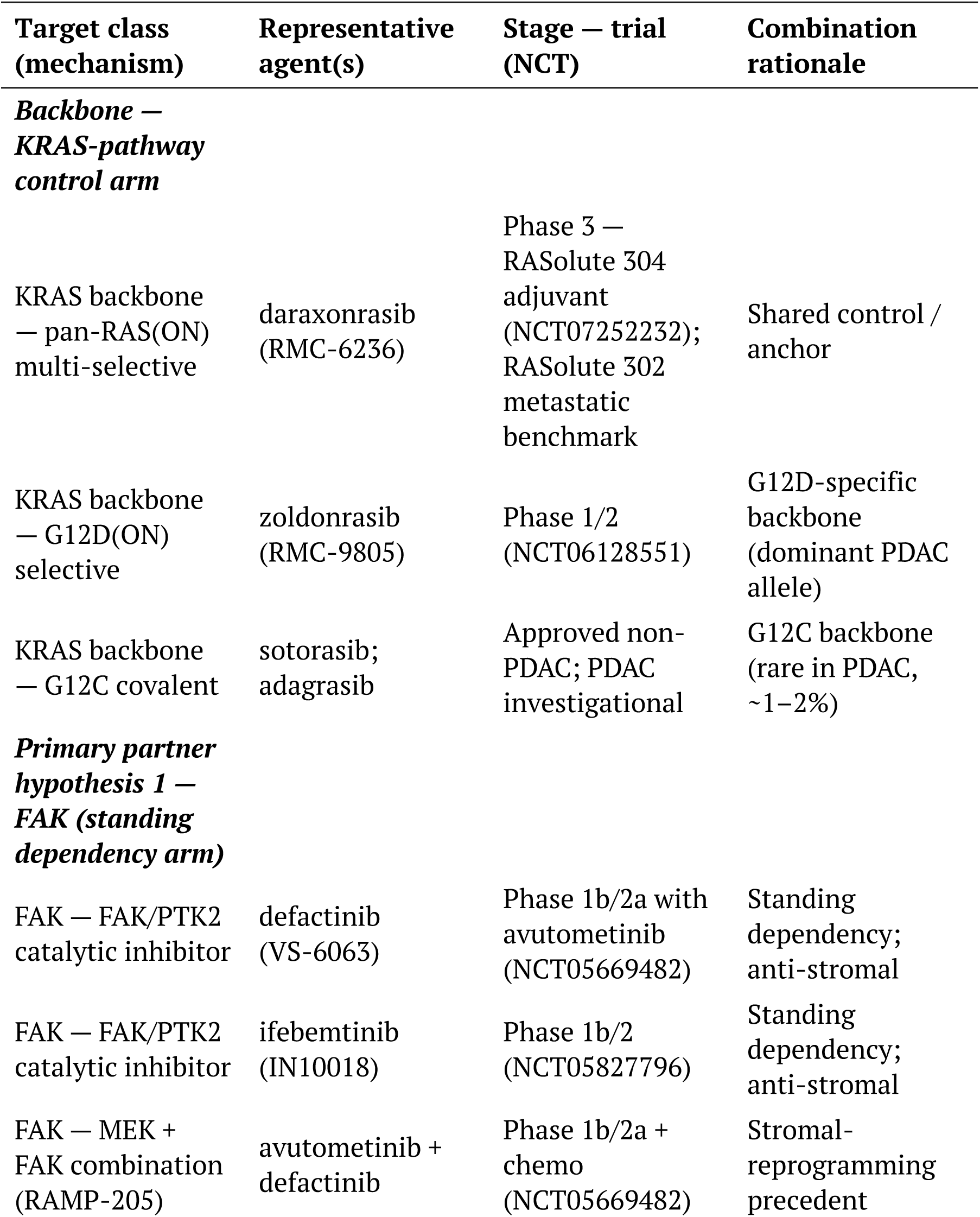
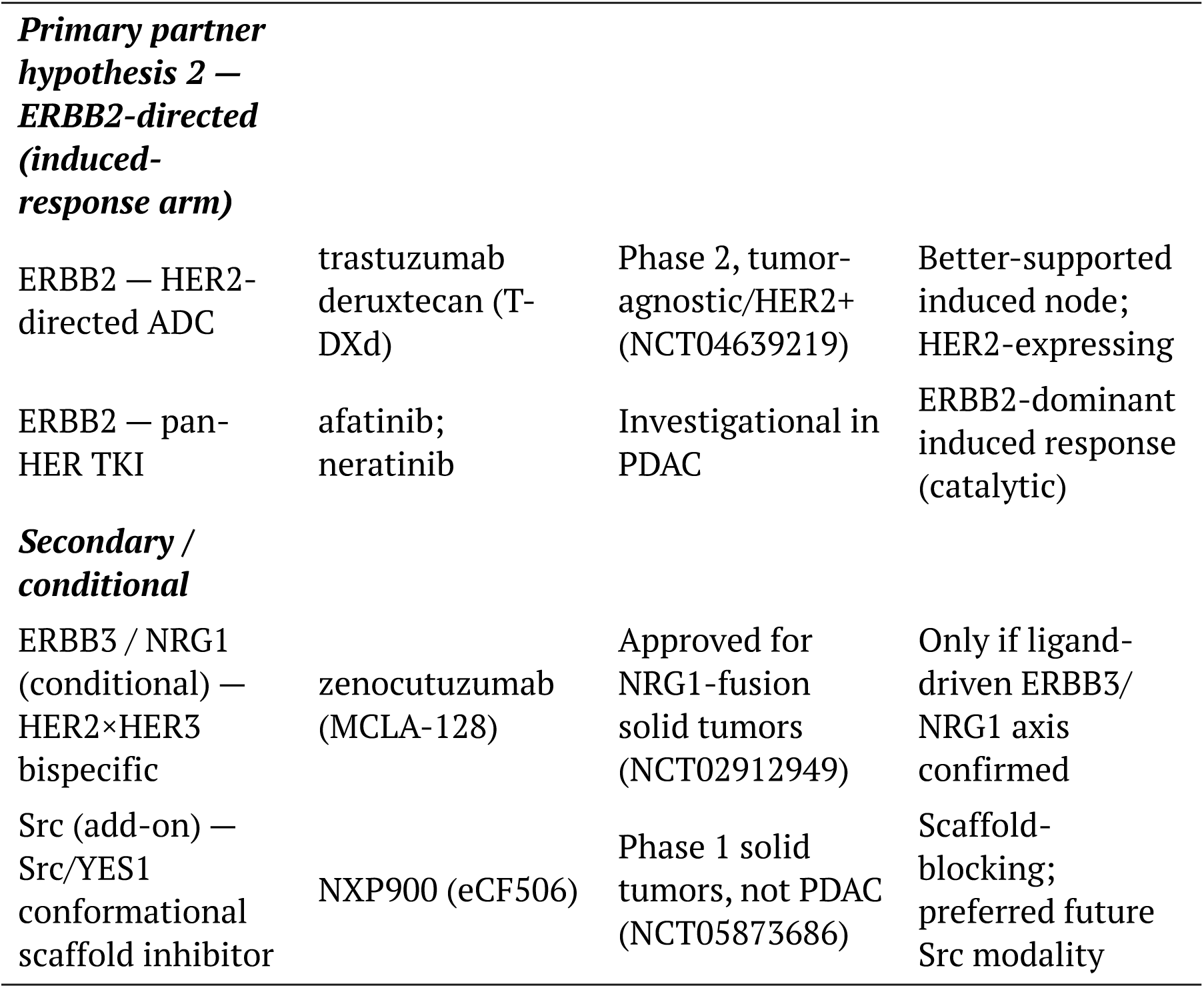
Candidate agents by target class for a KRAS-anchored combination platform in PDAC. Representative agents, modality, and PDAC development stage for each target class identified in this study. Agents are grouped by evidence tier — the KRAS backbone (shared control), the two primary partner hypotheses (FAK standing dependency; ERBB2-directed induced response, listed ahead of the weaker ERBB3/NRG1 axis, which is flagged conditional), and secondary/add-on concepts. Trial NCT identifiers are representative anchors verified against ClinicalTrials.gov; the table is a landscape reference, not an endorsement of any specific agent or regimen, and inclusion does not imply the combination has been tested. The historical negative single-agent Src trials (dasatinib, saracatinib) are catalogued in Supplementary Table S30 rather than here, as they inform the reinterpretation but are not partner candidates.

### A selection biomarker for the FAK arm

Because FAK dependency is a standing vulnerability rather than a KRAS-linked one, the CRISPR gene-effect score that nominates it (PTK2 Chronos) is not a clinically deployable stratifier, and the FAK arm needs a measurable enrichment marker of its own. We suggest a pragmatic hierarchy, in rough order of tissue-assay feasibility: (i) activated FAK by phospho-FAK (pFAK Tyr397) immunohistochemistry; (ii) a focal-adhesion/ invasion transcriptional signature or a CAF/fibrosis (desmoplasia) score read from bulk or spatial transcriptomics; (iii) the basal-like/squamous molecular subtype and stromal-high classifications, which are enriched for the adhesion-and-invasion biology FAK serves; (iv) an immune-exclusion readout (low intratumoral CD8), given FAK inhibition’s stroma-remodeling, T-cell–infiltration mechanism; and (v) ex vivo organoid or patient-derived-model FAK-inhibitor sensitivity as an integrated functional test. None of these is validated as a predictive biomarker in PDAC; each is a candidate stratifier that would itself require qualification against the FAK-combination readout, ideally co-developed within the platform’s biomarker-stratified design. The ERBB2 arm, by contrast, has a more conventional path — HER2 protein/amplification status, and where relevant an NRG1-fusion or ligand-axis readout for the conditional ERBB3 stratum.

### Required validation before any clinical inference

The central limitation is that every nomination here is computational and, in several places, underpowered; the claims are hypotheses about targets, not demonstrations of mechanism or efficacy. Before a trial could be justified, the minimum experimental package — set out concretely in **Box 1**, one row per nomination with a pre-stated readout and an explicit confirm/refute criterion — is: a KRAS-inhibitor time course in PDAC models; direct measurement of ERBB2/3 protein and phosphoprotein induction (and downstream pAKT/pERK recovery) after KRAS inhibition, since transcript up-regulation does not establish functional escape; combination-response testing of KRAS + ERBB and KRAS + FAK inhibition; and confirmation in organoid and stroma-inclusive or in vivo models rather than established cell lines alone. Two further caveats bound the interpretation. The ERBB escape evidence rests on cell-line transcriptomes at limited timepoints with small n and weak individual significance. And the network and GeneTerrain analyses are hypothesis-generating: network analysis places Src as an invasion-associated hub but does not support a specific Src–KRAS topological coupling (the shared-interactor overlap is no larger than a degree-matched null), and the GeneTerrain prognostic association requires external-cohort validation with adjustment for subtype, purity, stage, and stromal content before it can be read as more than corroborative. Elements of the clinical rationale drawn from conference abstracts should also be checked against their peer-reviewed form. What the data do support is narrower and, we believe, useful: a transparent, public-data nomination of two separable candidate KRAS-combination target classes, and a design by which each could be tested — once the validating experiments have been done.

*The trial concept above is a forward-looking hypothesis, not a validated protocol or a recommendation for current practice; the KRAS(ON) inhibitors it invokes are investigational, and every element requires prospective experimental and clinical testing*.

#### Box 1

**| Minimum experimental package to test the two nominations.**

Each computational nomination in this study maps to a specific, near-term experiment with a pre-stated readout and an explicit result that would confirm or refute it. None requires a clinical trial; all are feasible in a standard translational-PDAC laboratory. This is the bridge from the public-data hypotheses here to the evidence a trial would need.

| # • Nomination | Minimum experiment | Readout | Confirm / refute criterion |
| --- | --- | --- | --- |
| 1. KRAS inhibition induces an ERBB2-dominant adaptive response (transcript-level, underpowered) | KRAS-G12D/pan-RAS inhibitor time course (0–72 h) in ≥5 KRAS-mutant PDAC lines + organoids | ERBB2/ERBB3 **protein and phospho-protein**; pAKT/pERK recovery kinetics | ✓ ERBB2/3 and re-phosphorylation rise after MAPK suppression, protein-level, in a majority of models; ✗ Induction is transcript-only with no protein/phospho recovery, or absent in organoids |
| 2. ERBB inhibition is a productive KRAS-combination partner | KRAS inhibitor ± ERBB inhibitor (e.g. pertuzumab + trastuzumab, or a pan-HER TKI) combination-response matrix | Bliss/Loewe synergy; regrowth delay vs KRAS-inhibitor alone | ✓ Combination is synergistic and delays regrowth beyond monotherapy; ✗ No synergy; regrowth kinetics unchanged |

| # · Nomination | Minimum experiment | Readout | Confirm / refute criterion |
| --- | --- | --- | --- |
| 3. FAK is a standing (KRAS-independent) dependency | Genetic (sgPTK2) and pharmacological FAK inhibition at baseline, across KRAS-high- and KRAS-low-dependency lines | Growth/viability; confirm effect independent of KRAS-dependency magnitude | ✓ FAK loss impairs growth at baseline, comparably across the KRAS-dependency range; ✗ FAK effect tracks KRAS-dependency (i.e. not standing) or is absent |
| 4. FAK inhibition adds to a KRAS backbone, especially in stroma | KRAS inhibitor ± FAK inhibitor in stroma-inclusive / co-culture and in vivo (syngeneic or PDX) models | Tumor growth; stromal/CAF and immune-infiltrate markers | ✓ Added benefit over KRAS-inhibitor alone, with stromal remodeling; ✗ No added benefit in stroma-inclusive or in vivo settings |
| 5. The two programs are separable (FAK standing vs ERBB induced) | Head-to-head in the same models: does FAK dependency change on KRAS inhibition? does ERBB induction occur without FAK induction? | Paired ΔFAK-dependency and ΔERBB after KRAS inhibition | ✓ FAK dependency is present at baseline and unchanged, while ERBB rises only on KRAS inhibition; ✗ FAK is itself induced by KRAS inhibition (programs not separable) |
| 6. The Src-organized program is prognostic beyond generic EMT/stroma (single-cohort, exploratory) | External-cohort survival replication (CPTAC-3, Puleo, PRINCE) with adjustment for subtype, purity, stage, stroma | Multivariable Cox HR for the Src-neighborhood program | ✓ Program stays prognostic after adjustment in an independent cohort; ✗ Association vanishes under adjustment or does not replicate |
| A positive result on rows 1–2 would support the induced-ERBB arm; rows 3–4 the standing-FAK arm; row 5 tests the separability claim that motivates a multi-arm rather than single-combination design; row 6 tests the prognostic reading. A trial is justified only after the arm-specific rows return positive. |  |  |  |

## STAR Methods

### Data sources

All analyses used public data: cBioPortal (six PDAC mutation cohorts; TCGA-PAAD survival);^22^ DepMap (Chronos CRISPR gene effect, via the Open Targets Platform^21^ and DepMap 24Q4); GTEx (normal-tissue expression);^23^ STRING v12 (protein-protein interactome for the WINNER network-centrality ranking); ClinicalTrials.gov (trial records); ChEMBL (Src mechanism and agents); PubMed (literature); UCSC Xena Toil (TCGA-PAAD versus GTEx pancreas recompute); and GEO/ArrayExpress cohorts GSE240232 (genetic KRAS extinction), GSE303051 (pharmacological KRAS-G12C/D inhibition), GSE71729 (Moffitt), GSE154778 (Lin), CRA001160 (Peng), and E-MTAB-6134 (Puleo). Analysis used Python (pandas, scipy, lifelines, statsmodels).

### Perturbation analyses (candidate KRAS-inhibition adaptive response)

Two independent perturbation systems were analyzed. **Genetic extinction:** RNA-seq from three inducible-KRAS PDAC lines (GSE240232) was summarized as mean log₂ fold-change per gene panel under KRAS-ON versus KRAS-OFF, with per-panel paired t-tests across the three lines. **Pharmacological inhibition:** processed gene-level counts for 30 samples from GSE303051 — five PDAC lines (Pa01C, Pa03C, Pa14C, HPAC, MIA PaCa-2), each with three vehicle and three KRAS-G12C/G12D-inhibitor (MRTX-series) replicates — were converted to counts-per-million; per-line log₂ fold-change (drug versus vehicle) was computed for each gene, and a paired one-sample t-test across the five lines tested each gene panel. On-target pathway shutdown was verified via the direct MAPK-output genes DUSP6 and SPRY2. The steady-state co-dependency test correlated SRC, YAP1, and WWTR1 gene effect against KRAS gene effect (Pearson and Spearman) across 48 pancreatic lines.

### Genomic and functional dependency analysis

Somatic mutation and copy-number frequencies for SRC and KRAS were taken from six cBioPortal PDAC cohorts. DepMap Chronos gene effects were analyzed for pancreatic cell lines; dependency was defined at Chronos < −1 (primary) and < −0.5 (network ranking and threshold-robustness). The paired SRC/KRAS comparison used 48 unique pancreatic-proper lines (hepatopancreatic-ampulla lines excluded; lines deduplicated). The network dependency ranking scored each node by the fraction of pancreatic lines with gene effect < −0.5 and, continuously, by mean gene effect; the ranked set was the pre-specified KRAS/ Src/RTK/adhesion network, and the rank is reported as within-network, not genome-wide. Bootstrap 95% confidence intervals for dependency percentages and mean gene effects were computed by resampling cell lines with replacement (10,000 iterations). Normal-tissue expression used GTEx median TPM. **FAK KRAS-specificity:** per-line Chronos gene effect for PTK2 and KRAS was taken from the DepMap 24Q4 public CRISPRGeneEffect matrix for pancreatic-adenocarcinoma models (n = 45 with both genes); FAK–KRAS association was assessed by Pearson and Spearman correlation (bootstrap 95% CI, 10,000 iterations) and by comparing FAK gene effect and FAK-dependent fraction between the more- and less-KRAS-dependent halves of the cohort (median split, Mann–Whitney).

### Network-centrality ranking (WINNER)

As an orthogonal, dependency-independent prioritization, network centrality was computed with the WINNER algorithm^25^ (personalized-PageRank node scoring over a weighted protein-interaction graph; https://github.com/aimed-lab/WINNER). The KRAS/Src/RTK/adhesion module was used as seed nodes and expanded on a STRING v12 high-confidence interactome (combined score ≥ 700) to a 50-gene ranked universe with edge weights taken from STRING confidence scores. WINNER scores were computed at the default damping (σ = 0.85, 100 iterations); statistical significance of each node’s rank was assessed against 10,000 degree-preserving random networks (edge-swap null), yielding a per-node ranking p-value (the empirical fraction of random scores ≥ the observed score). FAK/PTK2 ranked 4th of 50 (ranking p = 0.0016). The full ranking is provided in the supplementary data.

### Survival analysis

Overall survival was modeled by continuous multivariable Cox regression (per–standard-deviation hazard ratio) in TCGA-PAAD, CPTAC-3, Puleo, and PRINCE, adjusted for age, stage, grade, and sex, with a DerSimonian-Laird random-effects meta-analysis across cohorts. The continuous per-SD hazard ratio was the pre-specified primary estimator; median-split and p-minimizing optimal-cutpoint models were computed only to demonstrate the cutpoint artifact, quantified by 1,000 permutations of survival against expression in TCGA-PAAD.^17,18^ Multiplicity across the tested gene family was controlled by Benjamini-Hochberg FDR; reporting follows prognostic-marker standards.^19,20^

### Differential expression and GeneTerrain analysis

Tumor-versus-normal effect sizes (Hedges’ g) for the originally ranked genes were computed in the Moffitt cohort before and after a leave-one-out proliferation adjustment. For GeneTerrain, TCGA-PAAD expression, clinical, and survival tables were assembled (178 primary tumors for outcome analysis; four solid-normal samples as the z-score baseline). A 42-gene SRC-centered module (SRC-pathway core, EMT/invasion, CAF/fibrosis, immune-evasion, epithelial identity) was converted to normal-baseline per-gene z-scores, from which module and composite terrain scores were computed on fixed coordinates. Prognostic analysis used Cox regression and Kaplan-Meier log-rank tests for the composite score, the SRC-pathway core score, and each module gene on overall survival and progression-free interval. Analysis used the GeneTerrain pipeline (Systems Pharmacology AI Research Center, UAB). **EMT/stromal confounder check (proxy analysis).** As an independent test of whether the prognostic signal is reducible to generic mesenchymal/stromal biology, the TCGA-PAAD RSEM expression matrix (177 tumors) was retrieved from cBioPortal and used to build two z-scored mean signatures: a 10-gene Src-neighborhood proxy (SRC, PTK2, EGFR, MET, ITGB1, ITGA2, VCL, MMP2, MMP14, KRT19) and a 14-gene EMT/stromal signature (VIM, FN1, ZEB1, ZEB2, SNAI1, SNAI2, TWIST1, CDH2, ACTA2, PDGFRB, COL1A1, COL1A2, COL3A1, FAP; epithelial markers excluded so all members share direction). Each was standardized and entered into univariate and joint per-SD Cox models for overall survival (176 patients, 92 events). This is a proxy for the Src-organized program, not a re-computation of the 42-gene GeneTerrain module, and is reported as a confounder check rather than a validation.

### Trial and network analysis

Src-agent trials were retrieved from ClinicalTrials.gov and classified by agent, phase, status, disease setting, and endpoint; agents and maximum clinical phase came from ChEMBL. The projected multi-arm survival curves are Weibull models anchored to the RASolute 302 daraxonrasib benchmark (Arm A) with additive combination effects assumed for the partner arms — illustrative of trial structure, not a prediction of effect size. Network integration (STRING/HAPPI interactomes, SPINNER edge re-weighting, degree-preserving configuration-model nulls) tested Src centrality and the SRC–KRAS shared-interactor overlap; the shared-interactor overlap was no larger than a degree-matched null, and is reported as module composition rather than coupling evidence. Figures were generated in matplotlib at 300 dpi; gene symbols are italicized, and effect sizes are reported with the test used.

## Data and code availability

All source data are public (accessions above). Every reported value is regenerated from a public data source. Reproduction scripts, per-figure source tables, and the multi-sheet supplementary data package accompany this manuscript; a provenance map records the data source underlying each section and figure. The essential public inputs, derived result tables, the standalone GeneTerrain reanalysis harness, the figures, and the manuscript are openly available at https://github.com/aimed-lab/kras-combo-therapy.

## Author contributions

J.Y.C. conceived the project, designed the approach, and implemented the data-analysis workflow. E.S. contributed the GeneTerrain analysis. G.O. and N.K. performed analysis support. Z.S. provided the computational-infrastructure support essential for completing the work. All authors reviewed and approved the manuscript.

## Acknowledgements

This work was supported in part by the University of Alabama at Birmingham through the Systems Pharmacology AI Research Center (SPARC). The authors gratefully acknowledge partial support from the following grants: NIH U54 OD036472 (CONNECT: Collaborative Network for Nurturing Ecosystems of Common Fund Team Science; Contact MPI/PI: Jake Y. Chen, 2024–2029); NIH OT2 OD032742 (Bridge2AI — Building an Interpretable Genomic Translator Using Maps of Cell Architecture; MPI, PI: Trey Ideker, 2022–2026); and the NAIRR Pilot (Computing for AI-enabled Systems Pharmacology and Drug Discovery; PI: Jake Y. Chen, 2025–2026). The content is solely the responsibility of the authors and does not necessarily represent the official views of the funding agencies.

## References

1. Sung H, et al. Global cancer statistics 2024: GLOBOCAN estimates of incidence and mortality worldwide for 34 cancers in 186 countries. CA Cancer J Clin 2026. PMID: 42417444.

2. Bailey P et al. Genomic analyses identify molecular subtypes of pancreatic cancer. Nature 2016. PMID: 26909576.

3. Yeatman TJ, et al. A renaissance for SRC. Nat Rev Cancer 2004. PMID: 15170449.

4. Nagaraj NS et al. Targeted inhibition of SRC kinase signaling attenuates pancreatic tumorigenesis. Mol Cancer Ther 2010. PMID: 20682659.

5. Poh AR et al. Functional roles of SRC signaling in pancreatic cancer: Recent insights provide novel therapeutic opportunities. Oncogene 2023. PMID: 37120696.

6. Evans TRJ et al. Phase 2 placebo-controlled, double-blind trial of dasatinib added to gemcitabine for patients with locally-advanced pancreatic cancer. Ann Oncol 2017. PMID: 27998964.

7. Renouf DJ et al. A phase I/II study of the Src inhibitor saracatinib (AZD0530) in combination with gemcitabine in advanced pancreatic cancer. Invest New Drugs 2012. PMID: 21170669.

8. Kelber JA et al. KRas induces a Src/PEAK1/ErbB2 kinase amplification loop that drives metastatic growth and therapy resistance in pancreatic cancer. Cancer Res 2012. PMID: 22589274.

9. Rozengurt E, et al. Crosstalk between KRAS, SRC and YAP Signaling in Pancreatic Cancer. Cancers (Basel) 2021. PMID: 34680275.

10. Han J et al. Stromal-derived NRG1 enables oncogenic KRAS bypass in pancreas cancer. Genes Dev 2023. PMID: 37775182.

11. Parkin A et al. Targeting the complexity of Src signalling in the tumour microenvironment of pancreatic cancer. FEBS J 2019. PMID: 31330086.

12. Jiang H et al. Targeting focal adhesion kinase renders pancreatic cancers responsive to checkpoint immunotherapy. Nat Med 2016. PMID: 27376576. doi:10.1038/nm.4123.

13. Liu X et al. Stromal reprogramming overcomes resistance to RAS-MAPK inhibition to improve pancreas cancer responses to cytotoxic and immune therapy. Sci Transl Med 2024. PMID: 39441902.

14. Wolpin BM et al. Daraxonrasib in Previously Treated Advanced RAS-Mutated Pancreatic Cancer. N Engl J Med 2026. PMID: 42090791. doi:10.1056/NEJMoa2505783.

15. Strickler JH et al. Sotorasib in KRAS p.G12C-Mutated Advanced Pancreatic Cancer. N Engl J Med 2023. PMID: 36546651.

16. Golan T et al. Maintenance Olaparib for Germline BRCA-Mutated Metastatic Pancreatic Cancer. N Engl J Med 2019. PMID: 31157963.

17. Altman DG et al. Dangers of using “optimal” cutpoints in the evaluation of prognostic factors. J Natl Cancer Inst 1994. PMID: 8182763.

18. Royston P et al. Dichotomizing continuous predictors in multiple regression: a bad idea. Stat Med 2006. PMID: 16217841.

19. Collins GS et al. Transparent reporting of a multivariable prediction model for individual prognosis or diagnosis (TRIPOD). BMJ 2015. PMID: 25569120.

20. Sauerbrei W et al. Reporting Recommendations for Tumor Marker Prognostic Studies (REMARK). J Natl Cancer Inst 2018. PMID: 29873743.

21. Ochoa D et al. The next-generation Open Targets Platform: reimagined, redesigned, rebuilt. Nucleic Acids Res 2023. PMID: 36399499.

22. Cerami E et al. The cBio cancer genomics portal. Cancer Discov 2012. PMID: 22588877.

23. GTEx Consortium, et al. The GTEx Consortium atlas of genetic regulatory effects across human tissues. Science 2020. PMID: 32913098.

24. Zhang A, Chen JY. AI-driven network biology identifies SRC as a therapeutic target in metastatic pancreatic adenocarcinoma. Intelligent Oncology 2025;1(3):233–243. doi:10.1016/j.intonc.2025.06.004.

25. Nguyen T, Yue Z, Slominski R, Welner R, Zhang J, Chen JY. WINNER: A network biology tool for biomolecular characterization and prioritization. Front Big Data 2022;5:1016606. PMID: 36407327. doi:10.3389/fdata.2022.1016606.

