## supplemental material for "Induced ERBB response and standing FAK dependency nominate separable KRAS-combination hypotheses in pancreatic cancer"

This document provides the detailed computational methods behind each supplementary table and figure, and the full legends. All analyses used publicly available data; no new experimental data were generated. Every result reported here is a computational nomination that requires experimental validation. The accompanying workbook V15_Supplementary_Data.xlsx (S0–S38) contains the underlying tabular data.

#### 1. Data sources and preprocessing

**cBioPortal (mutation and survival).** Somatic mutation frequencies for *KRAS* and *SRC* were retrieved for six PDAC cohorts (paad_cptac_2021, paad_icgc, paad_utsw_2015, paad_qcmg_uq_2016, paad_tcga_pan_can_atlas_2018, paad_tcga_gdc) through the cBioPortal API. TCGA-PAAD clinical/expression data were used for survival modelling. Frequencies are tabulated in **S5**.

**DepMap (CRISPR dependency).** Chronos gene-effect scores were obtained from DepMap 24Q4 and cross-referenced through the Open Targets Platform. Pancreatic lines were restricted to tissue = pancreas and deduplicated by cell line, yielding 48 unique lines (hepatopancreatic-ampulla lines excluded). A gene was called a dependency in a line at the standard Chronos threshold < −1; a relaxed threshold < −0.5 was used for the dependency-fraction ranking (Figure 3a). Per-line paired SRC/KRAS values are in **S6**; the broader gene panel in **S7**; the escape-panel genes in **S8**.

**GTEx (normal-tissue expression).** SRC median TPM across 54 GTEx v8 tissues (gencodeId ENSG00000197122.11) was used to assess the normal-tissue therapeutic window (Figure 5d).

**STRING v12 (interactome).** The high-confidence human protein–protein interaction network (combined score ≥ 700) was used to build the network for the WINNER centrality ranking (Section 4 below).

**GEO/ArrayExpress perturbation cohorts.** GSE240232 (genetic KRAS extinction), GSE303051 (pharmacological KRAS-G12C/D inhibition, 5 PDAC lines), GSE71729 (Moffitt), GSE154778 (Lin), CRA001160 (Peng), and E-MTAB-6134 (Puleo). GSE303051 raw per-sample gene-count files were downloaded, CPM-normalized, and collapsed to per-line log2 fold-change (treated vs vehicle) across the 5 lines.

#### 2. Perturbation analyses — the KRAS-inhibition escape state (S12, S17, S18, S19)

**Genetic extinction (S12).** In doxycycline-inducible KRAS-shutdown PDAC models, log2 fold-change on KRAS withdrawal was summarized per pathway panel (MAPK output, ERBB receptors, SFK, YAP/TAZ). MAPK output collapses on KRAS loss (mean log2FC −1.80, p < 0.001) while ERBB2 (+0.62, 3/3 lines, p = 0.007) and ERBB3 (+0.53, 3/3, p = 0.030) rise. FAK/PTK2 is not induced.

**Pharmacological inhibition (S17, S18).** For GSE303051, per-line log2FC on KRAS-G12C/D inhibition was computed for each gene. On-target MAPK-output genes fall sharply (DUSP6 −3.36, p = 0.004; SPRY2 −1.07, p = 0.062). ERBB2 (+0.58, 4/5 lines) and ERBB3 (+0.13, 4/5) rise directionally but with wide bootstrap CIs that cross zero (see Section 5), so the pharmacological escape signal is directionally consistent with the genetic system but underpowered at n = 5.

**Cross-system concordance (S19).** Genetic and pharmacological log2FC are placed side by side per gene with a direction-concordance flag. ERBB2, ERBB3, PTK2, WWTR1, and KRAS agree in direction across systems; SRC and YAP1 do not.

#### 3. Genomic and functional dependency analysis (S5, S6, S7, S13)

Mutation frequencies (S5) contrast *KRAS* (62–96%) with *SRC* (0–1.4%, no amplifications). The dependency landscape (S13, Figure 3a) ranks genes by the fraction of pancreatic lines in which knockout is lethal (Chronos < −0.5), computed on the 45 pancreatic-adenocarcinoma lines with complete DepMap 24Q4 CRISPR data — the single dependency universe used for every main-text dependency percentage (bootstrap 10,000 iterations for the confidence intervals): KRAS 93% (95% CI 84–100), FAK/PTK2 58% (42–71), TAZ/WWTR1 38% (24–53), EGFR 36%, ERBB2 22%, YAP1 20%, SRC 13% (4–24), ERBB3 9%, YES1 2%, FYN 0%. FAK is thus the strongest standing functional dependency after KRAS itself.

A separate 48-line paired subset (the lines with complete SRC/KRAS/YAP1/WWTR1 Chronos values) is used **only** for the baseline Src/KRAS co-dependency test (Figure 2c) and the S23 bootstrap on that co-dependency pair; it does not feed the Figure 3a ranking. This separation was made deliberately in this version so that all main-text dependency percentages come from one universe.

#### 4. Network-centrality ranking — WINNER (S22)

To answer “strongest among what universe,” we pre-specified a 50-gene module spanning the KRAS effector pathway, Src-family kinases, receptor tyrosine kinases, and focal-adhesion/integrin adhesion genes. The module was expanded through STRING v12 interaction partners (combined score ≥ 700) to a 477-gene universe with 1,110 high-confidence edges. Global node degree was computed from STRING partner counts (e.g. PTK2 degree 423, KRAS 919, SRC 1451).

We ran WINNER (Nguyen et al., *Frontiers in Big Data* 2022; PMID 36407327; doi:10.3389/fdata.2022.1016606; the AIMED-Lab tool the authors requested), a personalized-PageRank prioritization over the weighted interactome. The 50 module genes were seeds; ranking significance was assessed against 10,000 degree-preserving random networks (random seed 42; max_expansions = 0). WINNER is the algorithmic engine underlying the AIMED-Lab SPINNER application.

**Result:** within this pre-specified network, SRC ranks #1 (ranking p = 0.0000), FN1 #2 (p = 0.95, **not** significant against the degree-matched null), EGFR #3 (p = 0.037), and **FAK/PTK2 #4 of 50 (ranking p = 0.0016)**; ERBB2 ranks #13 (p = 0.57) and ERBB3 #28 (p = 0.021). The contrast between FN1 (rank 2 by raw centrality but not significant) and FAK (rank 4 and significant) shows that raw connectivity alone does not nominate a node — significance against the null does. FAK is therefore distinctive in scoring highly on both independent criteria — functional dependency (Section 3) and null-corrected network centrality — whereas Src leads centrality only. The full ranking is in **S22**.

**SPINNER visualization (Supplementary Figure S-SPINNER).** The ranked network was visualized with SPINNER (Seeded Protein Interaction Network Neighborhood Expansion and Ranking; https://github.com/aimed-lab/SPINNER), the AIMED-Lab application layer that wraps the WINNER node-scoring and WIPER edge-ranking engines. The 50-gene seed set was scored with SPINNER’s WINNER engine (100 iterations); edges among the 50 genes were drawn from the STRING v12 network (combined score ≥ 400, 666 edges). Node size is scaled to WINNER centrality; node colour marks rank significance against the 10,000-network degree-preserving null (p < 0.05); edge width and colour follow SPINNER’s confidence scale (the DEMA distance-bounded layout was reproduced faithfully from the SPINNER source). The edge list and per-node WINNER ranking are provided as V15_SPINNER_edgelist.tsv and V15_Supp_SPINNER_winner_ranking.csv in the Supplementary Data, so the entire ranking is reproducible from public STRING data with the open-source SPINNER package.

#### 5. Bootstrap confidence intervals (S23, S24)

All bootstrap CIs use 10,000 resamples (random seed 42 for dependency, 7 for the pharmacologic set) with percentile intervals.

**Dependency fractions and mean Chronos (S23), 48 lines.** KRAS: dependent (< −1) in 83.3% [72.9, 93.8], mean Chronos −2.14 [−2.41, −1.87]. SRC: 0.0% [0, 0] dependent, mean −0.22 [−0.29, −0.15]. YAP1 and WWTR1 intermediate. The KRAS/SRC separation (Cliff’s δ = +0.97) is unambiguous.

**Pharmacologic log2FC (S24), GSE303051 5 lines.** DUSP6 −3.36 [−4.23, −2.27] and WWTR1 −0.62 [−0.96, −0.29] have CIs excluding zero; **ERBB2 +0.58 [−0.08, 1.23] and ERBB3 +0.13 [−0.64, 0.81] both cross zero**, making explicit that the pharmacological ERBB signal indicates direction, not effect magnitude, at this sample size.

#### 6. Survival analysis (S9, S10, S11)

**Four-cohort meta-analysis (S9).** Continuous per-SD multivariable Cox HRs for SRC expression across TCGA-PAAD, CPTAC-3, Puleo, and PRINCE were pooled by random-effects meta-analysis: pooled HR 0.89 (95% CI 0.69–1.14), I² = 75.6%, Cochran’s Q p = 0.007 — no consistent prognostic effect.

**Cutpoint sensitivity (S10).** The reported “high Src → worse survival” association (HR ≈ 1.45) reproduces only under a p-minimizing “optimal” cutpoint. Permutation of survival against expression (1,000 permutations) shows this cutpoint search reaches p < 0.05 in 38% of permutations, giving a permutation-corrected p = 0.12.

**Leave-one-out proliferation adjustment (S11).** The differential-expression ranking that originally nominated Src is proliferation-driven: under LOO proliferation adjustment in the Moffitt cohort, Src collapses toward zero (Hedges’ g 0.20 → −0.09) alongside proliferation hubs RRM2/TOP2A/MKI67, while only KRAS retains a significant residual.

#### 7. GeneTerrain analysis (Figure 4 / main text)

A 42-gene SRC-centered module was projected to fixed 2D coordinates from a correlation-curated network over 178 TCGA-PAAD primary tumors (4 normal baselines for z-scoring), and the resulting GeneTerrain surfaces were compared between SRC-high and SRC-low tumors. Module-level scores were tested for survival association by univariate Cox. This is an internal within-cohort association (potentially confounded by stromal content, tumor purity, EMT state, and molecular subtype), not independent replication, and is the most exploratory analysis in the paper.

**Standalone reproduction harness (S-GeneTerrain-repro).** Because a prior review raised that this analysis depended on an in-house pipeline, the entire GeneTerrain result was re-implemented as an open, dependency-listed Python harness (V15_GeneTerrain_reanalysis_harness.py) that runs end-to-end on public TCGA-PAAD data with **no call to GeneTerrain as a service**. The harness ships with the raw inputs (PAAD_gene_expression_data.csv, 183 samples × 20,531 genes; PAAD_clinical_data.csv; PAAD_survival_data.csv) and a checksummed MANIFEST.csv (SHA-256 for every input and output). Steps: (i) normal-baseline z-scoring of each gene against the solid-normal samples; (ii) construction of five module scores — SRC-pathway core (SRC, YES1, FYN, LYN, PTK2, PXN, VCL, ITGB1, ITGA2, EGFR, MET), EMT/invasion, CAF/fibrosis, immune-evasion, and epithelial identity — with a composite SRC activity score = mean of the four aggressive modules minus the epithelial module; (iii) quartile grouping into SRC-High/SRC-Low; (iv) Mann–Whitney group tests and Benjamini–Hochberg-corrected per-gene Cox on OS and PFI. Re-running the harness reproduces the headline signal: SRC-pathway module OS HR = 2.24 (p = 1.3×10⁻⁵), PFI HR = 2.01 (p = 2.5×10⁻⁵), and an FDR-significant per-gene axis led by MET, KRT19, ITGA2, MMP7, and VCL (V15_Supp_GT_module_score_cox.csv, V15_Supp_GT_top12_FDR_genes.csv, V15_Supp_GT_module_score_tests.csv). This converts GeneTerrain from a black-box service into a fully reproducible analysis on public data.

#### 8. Drug-agent table (S25, main-text Table 1)

Candidate combination agents were compiled by target class (FAK, ERBB, SHP2, KRAS) from ChEMBL mechanism data and ClinicalTrials.gov, each anchored to a representative NCT record verified through the ClinicalTrials.gov API.

#### 9. Trial projection (Figure 6 / S16)

Projected overall-survival curves are Weibull models with the control arm anchored to the RASolute 302 daraxonrasib benchmark (OS 13.2 vs 6.7 months, HR 0.40). Partner arms are drawn under an assumed additive effect and depict trial geometry, not predicted efficacy.

#### 10. Reference validation (S21)

All 25 references were validated against PubMed/CrossRef (first author, year, journal matched); PMIDs/DOIs are tabulated in **S21**.

### Supplementary Figure Legends

The main-text figures (Figures 1–7) are described in the manuscript. This section documents the supplementary/underlying data views contained in the workbook and the provenance of each main figure’s data.

**Figure 1 (graphical abstract).** Three-lane schematic: oncogenic KRAS driver → KRAS-pathway inhibition (genetic + pharmacological) → two mechanistically separable candidate combination targets (induced ERBB2/3 escape; standing FAK dependency), which are not co-regulated (ρ = −0.43, n.s.). Rendered in matplotlib; all claims are computational nominations requiring validation.

**Figure 2 (escape state).** (a) Genetic KRAS extinction, per-panel log2FC for measured genes only (KRAS, ERBB2, ERBB3, SRC, PTK2, YAP1, WWTR1). (b) Pharmacological KRAS-G12C/D inhibition, 5 lines: per-line points, bootstrap-CI whiskers, and mean diamond; ERBB2/3 CIs cross zero. (c) Baseline co-dependency across 48 lines (Spearman ρ = +0.32, p = 0.03) — positive, so not a standing trade-off. Data: S18, S19, S23, S24, S6, S8.

**Figure 3 (dependency + trial setting).** (a) Fraction of the 45 pancreatic-adenocarcinoma DepMap 24Q4 lines dependent (Chronos < −0.5) per gene — the single dependency universe (Section 3): KRAS 93%, FAK/PTK2 58%, TAZ/WWTR1 38%, EGFR 36%, ERBB2 22%, YAP1 20%, SRC 13%, ERBB3 9%, YES1 2%, FYN 0%. FAK 58% is the strongest after KRAS. Inset: per-line FAK/PTK2 vs KRAS gene effect in the same 45 lines, Spearman ρ = +0.05, n.s. (b) Efficacy-powered PDAC Src trials by disease setting: 4 trials (N=316) advanced/metastatic vs 1 (N=8) resected/adjuvant. (c) Separability summary: FAK standing/baseline vs ERBB2/3 induced. Data: S13, S4, S26.

**Figure 4 (GeneTerrain — promoted 3-panel main figure).** (a) SRC-High − SRC-Low GeneTerrain difference map: the 42 module genes at fixed 2-D coordinates, coloured by the mean normal-baseline z-score difference between top- and bottom-quartile SRC-activity tumors; the invasion/EMT/CAF cluster (SNAI2, ZEB1/2, VIM, FN1, collagens, FAP, MMP2) is coordinately elevated (warm) while epithelial-identity genes (EPCAM, CDH1, keratins) are depleted (cool). (b) Per-SD OS hazard ratios for the Src-neighborhood score vs an EMT/stromal signature, univariate and mutually adjusted (n = 176, 92 events); the Src-neighborhood program retains HR 1.96 after EMT adjustment while EMT collapses to 0.90. (c) Real per-patient Kaplan–Meier separation at the median Src-neighborhood score (log-rank p = 6×10⁻⁴, per-SD HR 1.87, C = 0.63). The full dashboard is Supplementary Figure S-GeneTerrain and the analysis is reproducible from public data (Supplementary Figure S-GeneTerrain-repro). Real-data figure; exploratory single-cohort association. Data: main text, S27, S28.

**Figure 5 (Src molecular profile).** (a) KRAS vs SRC mutation frequency, six cohorts. (b,c) KRAS vs SRC CRISPR dependency, 48 lines. (d) SRC normal-tissue expression, GTEx. Data: S5, S6, GTEx.

**Figure 6 (Src prognostic reinterpretation).** (a) Four-cohort per-SD SRC survival forest, pooled HR 0.89. (b) Cutpoint permutation null (38% reach p<0.05; corrected p=0.12). (c) LOO proliferation adjustment. Data: S9, S10, S11.

**Figure 7 (platform-trial concept).** Multi-arm platform schematic: biomarker-stratified resected/adjuvant KRAS-mutant PDAC randomized to a shared KRAS-inhibitor control (Arm A) and separable KRAS + FAK (Arm B) and KRAS + ERBB (Arm C) partner arms. Illustrative projected survival curves for this design are shown in Supplementary Figure S-proj (geometry only). Data: schematic.

#### Supplementary figures (added in V11–V12)

**Supplementary Figure S-GeneTerrain (full GeneTerrain dashboard).** The complete multi-panel GeneTerrain analysis across 178 TCGA-PAAD primary tumors (moved here from the V11 main text so the main Figure 4 carries only the single takeaway). Panels: (A) GeneTerrain workflow; (B) SRC-centered 42-gene network topology; (C–E) SRC-low, SRC-high, and difference terrain surfaces (elevated EMT, CAF/fibrosis, immune-evasion ridges in SRC-high); (F) module-level biological separation by patient group; (G) Cox survival associations of GeneTerrain scores (SRC-pathway core per-score OS HR 1.96, 95% CI 1.44–2.65, p = 1.5×10⁻⁵; broad composite only weakly/non-robustly prognostic, HR 1.31, p = 0.037); (H) BH-FDR-corrected key prognostic genes (MET, KRT19, ITGA2, VCL, EGFR, MMP14, MMP7, SNAI2); (I) Kaplan-Meier separation (log-rank p = 0.011 OS, 0.0036 PFI); (J) per-SD OS HR 2.24, PFI HR 2.01. Single-cohort, investigator-curated module, four normal-baseline samples, in-house pipeline — internal corroboration requiring external validation, not independent replication. Data: main text.

**Supplementary Figure S-FAK (FAK KRAS-specificity).** Per-line CRISPR gene effect for FAK/PTK2 versus KRAS across 45 PDAC cell lines (DepMap 24Q4 public CRISPRGeneEffect). Left: scatter with the FAK–KRAS association (Pearson r = +0.09 p = 0.57; Spearman ρ = +0.05 p = 0.75; bootstrap 95% CI −0.25 to +0.35). Right: FAK gene effect and FAK-dependent fraction in the more- versus less-KRAS-dependent halves of the cohort (median split): FAK-dependent 57% vs 59%, Mann–Whitney p = 0.85. FAK dependency is not KRAS-specific — it is a standing vulnerability independent of KRAS-dependency magnitude. Data: S26.

**Supplementary Figure S-EMT (Src-neighborhood vs EMT/stromal signature).** TCGA-PAAD (176 tumors, 92 events, cBioPortal RSEM). A 10-gene Src-neighborhood proxy score and a 14-gene EMT/stromal signature were entered into per-SD Cox models. Univariate: Src-neighborhood OS HR 1.87 (1.42–2.46, p = 8×10⁻⁶), EMT HR 1.28 (1.03–1.59, p = 0.03). Joint model: Src-neighborhood retains HR 1.97 (1.45–2.66, p = 1×10⁻⁵) while EMT collapses to HR 0.90 (p = 0.45); concordance 0.63 vs 0.51. This is a proxy confounder check (not the 42-gene GeneTerrain module): the Src-organized program carries prognostic information a generic mesenchymal signature does not. Data: S27.

**Supplementary Figure S-proj (projected platform-trial survival).** Illustrative Weibull overall-survival curves for the multi-arm platform of Figure 7. Arm A (KRAS inhibitor alone) is anchored to the daraxonrasib RASolute 302 metastatic benchmark (OS 13.2 vs 6.7 months, HR 0.40); the partner arms B/C are drawn under an assumed additive effect. These depict trial geometry only and are not a prediction of efficacy. Data: S16.

**Supplementary Figure S-SPINNER (SPINNER/WINNER network prioritization).** The pre-specified 50-gene KRAS/Src/RTK/adhesion module rendered with the AIMED-Lab SPINNER application (WINNER node scoring, WIPER edge context; https://github.com/aimed-lab/SPINNER). Node size is proportional to WINNER centrality; nodes are coloured by rank significance against a 10,000-network degree-preserving null (red = p < 0.05: SRC #1, EGFR #3, FAK/PTK2 #4, ERBB3 #28, plus GRB2 and PTK2B); edges are the STRING v12 high-confidence interactions among the 50 genes (combined score ≥ 400, 666 edges), drawn with SPINNER’s confidence-scaled width and colour. FN1 ranks #2 by raw centrality but is not significant against the null (p = 0.95), whereas FAK’s rank-4 position is (p = 0.0016) — connectivity alone does not nominate a target. Reproducible from V15_SPINNER_edgelist.tsv with the open-source SPINNER package. Data: S22, V15_Supp_SPINNER_winner_ranking.csv.

**Supplementary Figure S-GeneTerrain-repro (open reproduction of the GeneTerrain prognostic result).** Prognostic forest from the standalone reproduction harness (Section 7), run entirely on public TCGA-PAAD data with no GeneTerrain service dependency. Module-level Cox hazard ratios reproduce the in-house pipeline: the SRC-pathway score is prognostic for poor outcome (OS HR 2.24, p = 1.3×10⁻⁵; PFI HR 2.01, p = 2.5×10⁻⁵), as are the EMT-invasion, CAF-fibrosis, and immune-evasion modules (all HR > 1). The epithelial-identity module score is also associated with worse outcome in this harness (OS HR 5.8, PFI HR 6.0), not protective: all six member genes point in the poor-prognosis direction (per-gene Cox OS HR > 1), and five reach significance (KRT19 1.96, KRT8 1.51, KRT18 1.49, CDH1 1.46, CLDN4 1.40; all p < 0.01), while EPCAM is directionally concordant but not individually significant (HR 1.19, p = 0.17). The module score inherits that overall direction. This is a property of keratin/epithelial-marker prognosis in this cohort, not an epithelial-protective effect; per-gene hazard ratios are in V15_Supp_GT_module_score_cox.csv and the harness per-gene Cox output. Top BH-FDR-significant per-gene associations (MET, KRT19, ITGA2, MMP7, VCL) reproduce the in-house pipeline. Inputs and outputs are SHA-256-checksummed in the harness MANIFEST.csv. Data: V15_Supp_GT_module_score_cox.csv, V15_Supp_GT_top12_FDR_genes.csv, V15_Supp_GT_module_score_tests.csv.
